# Design of highly functional genome editors by modeling the universe of CRISPR-Cas sequences

**DOI:** 10.1101/2024.04.22.590591

**Authors:** Jeffrey A. Ruffolo, Stephen Nayfach, Joseph Gallagher, Aadyot Bhatnagar, Joel Beazer, Riffat Hussain, Jordan Russ, Jennifer Yip, Emily Hill, Martin Pacesa, Alexander J. Meeske, Peter Cameron, Ali Madani

**Affiliations:** Profluent Bio, Berkeley, CA, USA; Laboratory of Protein Design and Immunoengineering, École Polytechnique Fédérale de Lausanne and Swiss Institute of Bioinformatics, Lausanne, Switzerland; Department of Microbiology, University of Washington, Seattle, WA, USA

**Keywords:** genome editing, large language models, protein design

## Abstract

Gene editing has the potential to solve fundamental challenges in agriculture, biotechnology, and human health. CRISPR-based gene editors derived from microbes, while powerful, often show significant functional tradeoffs when ported into non-native environments, such as human cells. Artificial intelligence (AI) enabled design provides a powerful alternative with potential to bypass evolutionary constraints and generate editors with optimal properties. Here, using large language models (LLMs) trained on biological diversity at scale, we demonstrate the first successful precision editing of the human genome with a programmable gene editor designed with AI. To achieve this goal, we curated a dataset of over one million CRISPR operons through systematic mining of 26 terabases of assembled genomes and meta-genomes. We demonstrate the capacity of our models by generating 4.8x the number of protein clusters across CRISPR-Cas families found in nature and tailoring single-guide RNA sequences for Cas9-like effector proteins. Several of the generated gene editors show comparable or improved activity and specificity relative to SpCas9, the prototypical gene editing effector, while being 400 mutations away in sequence. Finally, we demonstrate an AI-generated gene editor, denoted as OpenCRISPR-1, exhibits compatibility with base editing. We release OpenCRISPR-1 publicly to facilitate broad, ethical usage across research and commercial applications.

## Introduction

Genome editing technologies, including those derived from prokaryotic CRISPR-Cas systems, have revolutionized life science research and are poised to transform medicine and agriculture. Single-protein CRISPR-Cas effectors, including the widely-adopted Cas9 nuclease from Streptococcus pyogenes (SpCas9), have been utilized in biotechnology owing to their simplicity, robustness, and compact form. In order to diversify the CRISPR toolbox and expand editing capabilities, new systems have been mined across diverse microbial and viral genomes. While these novel systems have been sought for specific properties, such as small size or extended protein stability in biofluids (1, 2), they typically exhibit trade-offs in critical attributes such as basal activity in target cells, PAM selectivity, thermal optima, or in vitro biochemical properties, ultimately limiting their reach (2–5).

Repurposed CRISPR systems have been optimized for biotechnology using a range of protein engineering approaches, including directed evolution and structure-guided mutagenesis. Directed evolution of Cas proteins has proven extremely powerful yet can be limited by the rugged and non-convex nature of the fitness landscapes (6–9), along with the difficulty of implementing selection-based screening in human cells. Structure-guided rational mutagenesis offers an alternative or synergistic approach that has proven successful for improving Cas9 basal activity and specificity in human cells (10–13). Similar results may be achievable with structure-conditioned protein sequence design models (14, 15), which learn the mapping from structure to sequence from data. However, both of these approaches are dependent on explicit structural hypotheses, either in the form of mechanistic understanding for rational mutagenesis or solved structures representing key functional states for computational design, which are difficult to obtain for functions more complex than simple binding interactions.

Protein language models eschew explicit structural hypotheses and instead learn the co-evolutionary blueprint underlying protein function (16). When pre-trained on large sets of diverse protein sequences, language models learn to represent structure and function without supervision (17, 18). Subsequent fine-tuning of these models yields family-specific specialists that generate novel proteins adhering to the functional constraints of their family yet diverging significantly in sequence space. This approach has been validated through the design of functional lysozymes (19), with sequence identities as low as 31% to any known natural protein, and demonstrated in silico across multiple families (20). However, it remains to be seen how well this strategy performs for protein families with multiple complex functions, such as CRISPR-Cas effectors. For these systems, protein function is dependent on several external factors, including protein-specific accessory RNAs and target site preferences, which must be accounted for to achieve editing. Further, for therapeutic applications, gene editors must be functional in more challenging human cell environments, beyond the scope of most protein design efforts.

In this work, we demonstrate that language models can effectively generate diverse CRISPR-Cas proteins spanning a broad set of families. Moreover, we demonstrate that generated Type II effector proteins can assemble as functional gene editors in human cells, despite being hundreds of mutations away from any known natural protein. To our knowledge, this represents the first successful editing of the human genome by proteins designed entirely with machine learning. We perform extensive characterization of one exemplar editor, which we denote OpenCRISPR-1, and show that it is highly functional and specific. We release OpenCRISPR-1 for broad ethical usage across research and commercial applications.

## Results

### Language models generate diverse CRISPR-Cas proteins

Generative protein language models are typically pre-trained on large datasets of natural protein sequences spanning diverse phylogenies and functions (18). These models are capable of generating realistic protein sequences that reflect the distribution and properties of natural proteins (21). However, for specific applications, such as the generation of novel gene editors, it is necessary to steer generation towards particular subsets of protein families of interest (Fig. 1a).

**Figure 1.**
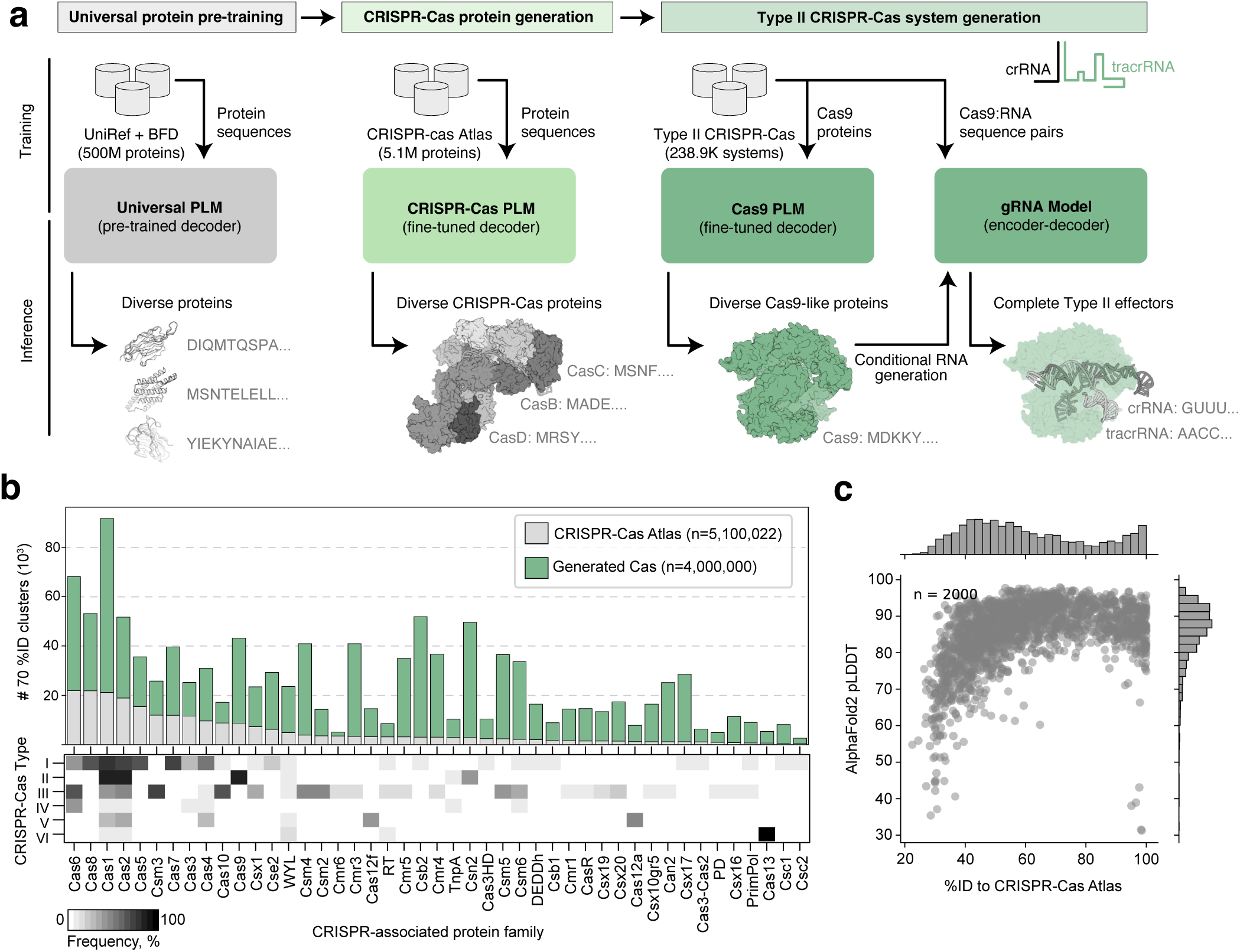
Generation of diverse CRISPR-associated protein families. a) Overview of the language modeling approach to design CRISPR-Cas systems. Language models learn the general constraints of protein evolution through pre-training on diverse proteins spanning the evolutionary tree, then are specialized for design by fine-tuning on CRISPR-associated protein and nucleic acid data. b) Expansion of the sequence diversity for 45 CRISPR-associated protein families, measured by the number of clusters (at 70% sequence identity) for natural proteins and novel clusters from generated sequences. Stacked bars are colored by the source of the sequences making up their clusters (CRISPR-Cas Atlas: recovered from CRISPR-Cas mining; Generated: four million generated proteins from this study). Heatmap indicates the frequency of each protein family across different CRISPR-Cas system types from the CRISPR-Cas Atlas. c) AlphaFold2 was used to predict structures for 2,000 randomly selected generated proteins. The scatterplot shows the distribution of mean pLDDT and the percent identity to natural proteins from the CRISPR-Cas Atlas.

To this end, we performed exhaustive data mining to construct, to our knowledge, the most extensive dataset of CRISPR operons curated to-date, including CRISPR-associated (Cas) proteins, CRISPR arrays, tracrRNAs, and protospacer adjacent motifs (PAMs) (Fig. S1). We refer to this resource as the CRISPR-Cas Atlas. Using a custom pipeline, we searched 26.2 terabases of assembled microbial genomes and metagenomes, spanning diverse phyla and biomes, to uncover 1,246,163 CRISPR-Cas operons, including over 389,470 single-effector systems classified as Type II, Type V, or Type VI (Fig. S1). Our resource displayed expanded natural diversity compared to curated databases like CRISPRCasDB and CasPDB (Fig. S1e), as well as UniProt, the world’s largest protein resource. Across all Cas families, the CRISPR-Cas Atlas has on average 2.7x more protein clusters than UniProt using a 70% identity clustering threshold, and even greater expansions for families like Cas9 (4.1x), Cas12a (6.7x), and Cas13 (7.1x).

Although CRISPR-Cas proteins carry out a variety of distinct roles, they often do so through a common set of domains and structural motifs (22). For example, Type II and V effector proteins typically contain a RuvC-like nuclease domain, which is responsible for cleavage of DNA. Similarly, both adaptor and effector proteins may have PAM-interacting domains, which are responsible for recognition of specific DNA sequence motifs. Given these commonalities, we reasoned that a single model for all CRISPR-Cas proteins would be capable of generating diverse sequences across families. To generate novel CRISPR-Cas proteins, we fine-tuned the ProGen2-base language model (18) on the CRISPR-Cas Atlas, balancing for protein family representation and sequence cluster size (Fig. 1a). From this model, we generated four million sequences; half were generated directly from the model, while the other half were prompted with up to 50 residues from the N- or C-terminus of a natural protein to guide generation towards a particular family. To assign the generated sequences to CRISPR-Cas families, we aligned each sequence to the CRISPR-Cas Atlas using BLAST (23), selecting the family of the best alignment for classification. All generated sequences were filtered according to a set of BLAST and HMM alignment criteria to remove degenerate sequences. To assess the novelty and diversity of generated sequences, we clustered the generated and natural sequences for each family at 70% identity using MMseqs2 (24). Overall, the generated sequences represented a 4.8-fold expansion of diversity compared to natural proteins from the CRISPR-Cas atlas (Fig. 1b). For families with few natural proteins, such as Cas13 and Cas12a, generated sequences represent an 8.4- and 6.2-fold increase in diversity, respectively. Among the sequences guided towards a particular family, we typically observed near-perfect adherence to the family of interest with fifty or fewer residues provided, suggesting that generation is steerable with minimal context.

Having successfully generated novel sequences for all targeted families, we next evaluated the rate of novel cluster generation by the model with respect to the number of sequences sampled. For each family, we counted the number of distinct clusters within subsets ranging from one sequence to the full set (Fig. S2). In general, the rate of novel cluster generation for generated sequences significantly outpaced that of natural sequences, suggesting we have not saturated the generative capacity of the models. Furthermore, our four million generations represent a finite sampling for only three days on 16 GPUs and the generative models are capable of generating vastly greater numbers of CRISPR-Cas sequences. Among the generated sequences for each family, we observe significant sequence diversity, with the median identity to the nearest natural protein typically falling between 40-60% (Fig. 1c). Despite this considerable deviation in sequence space, the generated proteins were confidently predicted by AlphaFold2 (with 81.65% of structures having mean pLDDT above 80) (Fig. 1c) and were predicted to adopt folds highly similar to natural proteins from the same family (Fig. S3), suggesting the generated sequences may be functional.

### Language models generate diverse Type II effectors

While many CRISPR-Cas proteins have been leveraged for genome editing (27, 28), Cas9 remains the most widely employed. To generate novel Cas9-like sequences, we prompted the CRISPR-Cas model with 50 residues from the N- or C-terminus of Cas9s sampled from the CRISPR-Cas Atlas. However, only 27.6% of these prompted generations survived our strict sequence viability filters. To more efficiently and accurately generate viable Cas9-like sequences, we fine-tuned another language model using only the 238,917 Cas9 sequences from the CRISPR-Cas Atlas (Fig. 1a and Fig. S1). This model produced viable Cas9-like sequences at twice the rate of the CRISPR-Cas model (54.2%, Fig. S4a) and did not require any prompting.

To explore the latent sequence distribution of Type II effectors, we used the Cas9 model to generate 1 million Cas9 proteins. The resulting viable generations (n=542,042) were clustered together with natural Cas9s at 40% identity and used as input to construct a maximum likelihood phylogenetic tree (Fig. 2a). Strikingly, the resulting landscape was dominated by generated proteins, which comprised 94.1% of the total phylogenetic diversity (as measured by cumulative branch length) and resulted in a 10.3-fold increase in diversity relative to the entire CRISPR-Cas Atlas (Fig. 2b). Novel phylogenetic groups were distributed across the tree, suggesting that the model has captured the full diversity of Cas9 and is not overfitting to any particular lineage. Generated sequences diverged considerably from the CRISPR-Cas Atlas, with an average identity of only 56.8% to any natural sequence (Fig. 2c). Although the number of generated proteins is large, the number of novel clusters does not appear to have saturated (Fig. S4b), suggesting that many more novel Cas9-like proteins could be generated. Next, we updated the phylogenetic tree with 79 diverse Cas9 orthologs that have been biochemically characterized (5) or used as genome editors (25). We observed considerable diversity of generated proteins in the vicinity of these characterized orthologs, suggesting that the model is capable of generating proteins with a variety of functional properties, including PAM specificity, temperature-dependent activity, DNA cleavage patterns, or high activity in human cells.

**Figure 2.**
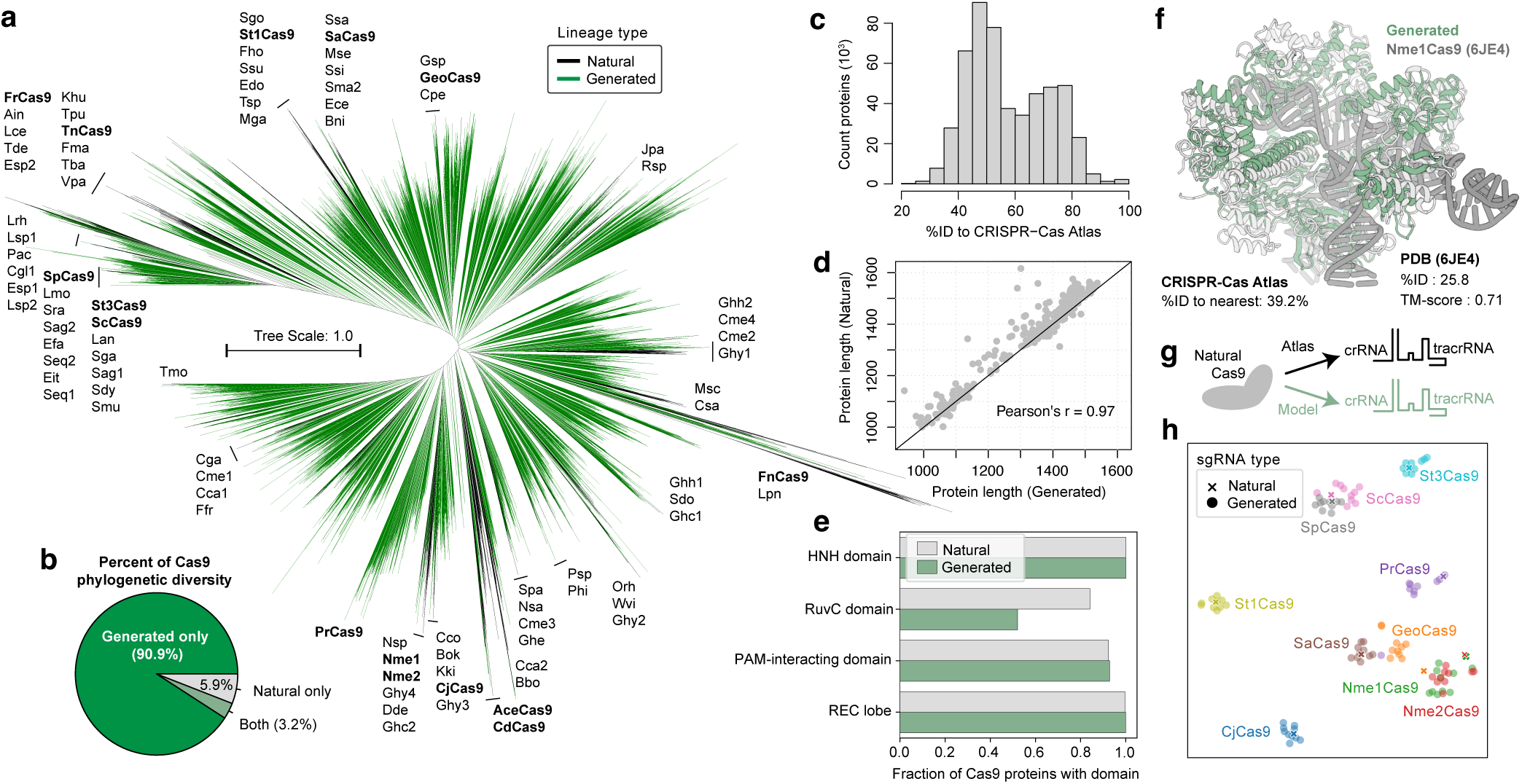
Language models generate complete Type II effector systems. a) Phylogenetic tree of natural and generated proteins clustered at 40% identity (n=15,340 cluster representatives). Biochemically characterized Cas9s from Gasiunas et al. (5) are labeled and Cas9 proteins used as genome editors are shown in bold (25). Lineages are colored black if they contain any natural protein while lineages are colored green if they are exclusively represented by generated proteins. b) Piechart indicates the percent of phylogenetic diversity represented by natural or generated proteins. Phylogenetic diversity was calculated as the cumulative branch length of subtrees represented by a given set of sequences. c) Distribution of the identity of generated Cas9 to the nearest protein in the CRISPR-Cas Atlas. d) Comparison of protein length between natural and generated proteins in the same 50% ID clusters. e) Fraction of generated and natural Cas9 proteins containing key functional domains according to structural searches with Foldseek against SCOPe families. In total, 79.2% and 48.2% of natural and generated proteins were functionally complete, respectively. f) Predicted structure for highly novel Cas9-like protein selected from a 30% identity cluster with 423 members composed entirely of generated sequences. Despite high sequence novelty (39.2% identity to CRISPR-Cas Atlas), the predicted structure bears strong structural resemblance to Nme1Cas9 (PDB ID 6JDQ, TM-score = 0.71). g) Naturally occurring and generated crRNAs and tracrRNAs were obtained for a set of ten effector proteins. h) sgRNAs were formed from RNA components and embedded into a two-dimensional space by t-SNE (26) according to the pairwise edit distances. Each point represents an sgRNA sequence with colors corresponding to source protein.

To further assess the viability of the generated proteins, we compared the sequence lengths of generated and natural sequences (Fig. 2d and Fig. S4c). Overall, generated sequences closely matched the length of natural proteins from the same protein cluster, with a Pearson correlation of 0.97 (Fig. 2d). To assess the structural viability of generated Cas9-like proteins, we used AlphaFold2 (29) to predict the structures of 5,000 generated and 5,000 natural sequences, selected from the largest protein clusters in each set. The vast majority of structures were predicted confidently (with 99.4% having mean pLDDT above 80), including for Cas9-like proteins with as low as 60% identity to any natural protein (Fig. S5a), and showed significant overlap with experimentally determined structures from the PDB (Fig. S5b). By aligning the structures against curated families from the SCOPe database (30), we revealed the presence of core Cas9 domains in most generated proteins and at a similar rate as naturals (Fig. 2e). This included the HNH and RuvC nuclease domains (100% and 52.1%, respectively), which are responsible for DNA cleavage, as well as the PAM-interacting domain (92.9%) and target recognition (REC) lobe (99.9%) (Fig. 2e). This structural and functional completeness extended to even the most novel proteins, including a subset that belonged to 30% identity clusters composed entirely of generated proteins. One such protein had a predicted structure resembling Nme1Cas9 (TM-score = 0.71) despite sharing only 25.8% identity to Nme1Cas9 and maximum of 39.2% identity to any protein in the CRISPR-Cas Atlas (Fig. 2f).

Beyond the effector protein, Type II systems are also dependent on a guide RNA (gRNA) that is required for target recognition and cleavage. The gRNA is composed of a targeting RNA sequence (spacer), CRISPR RNA (crRNA) repeat, and trans-activating CRISPR RNA (tracrRNA). The tracrRNA and crRNA components are typically derived from natural systems, while the spacer sequence is programmed to match the target DNA site for gene editing applications. For a given generated protein, we sought to model the crRNA and tracrRNA sequences that would enable effector function. From the CRISPR-Cas Atlas, we collected 112,212 Type II effector proteins for which we could confidently identify, orient, and align the corresponding crRNA and tracrRNA sequences. These data were used to train a sequence-to-sequence gRNA model that conditionally generates crRNA and tracrRNA sequences for a given protein (Fig. 1a). As an initial validation of the gRNA model, we designed ten gRNAs for a set of effector proteins previously used as genome editors (Fig. 2g). Each of the designed gRNAs, as well as the natural gRNAs from metagenome mining, were formatted into single-guide RNAs (sgRNAs) and embedding according to their pairwise edit distances with t-SNE (26). We observed that the model-designed sgRNAs were most similar to the naturally derived sgRNAs for each protein (Fig. 2h). As additional validation, we found crRNA:tracrRNA pairs often formed the canonical duplex (Fig. S6) and that the model could accurately predict the compatibility of sgRNAs between diverse Cas9 orthologs (Fig. S7). Together these results suggest that the model can be used to generate functional sgRNAs for novel generated Cas9-like proteins.

### Generated gene editors are functional in human cells

To generate Cas9-like proteins for experimental characterization, we employed a constrained generation strategy wherein we prompted the language model fine-tuned on Cas9 proteins using either the N-terminal segment or the C-terminal PAM-interacting domain (PID) of SpCas9 (Fig. S8). In doing so, we reasoned that the generated sequences would maintain compatibility with both the PAM and sgRNA preferences of SpCas9, facilitating direct comparison of protein activity across the same genomic targets with the same sgRNA. In total, we generated 200,000 and 150,000 Cas9-like proteins prompted by the N-terminal segments and C-terminal PIDs, respectively. From the SpCas9 PID-conditioned sequences, we selected 82 designs for characterization. To create fully generated effector proteins, we combined our generated N-terminal segments and PID domains. From these designs an additional set of 127 sequences was selected for characterization based on a combination of language model scores and auxiliary predictors of compatibility with SpCas9’s PAM and tracrRNA. All together, we selected 209 Cas9-like proteins for subsequent functional analysis in human cells (Fig. 3a).

**Figure 3.**
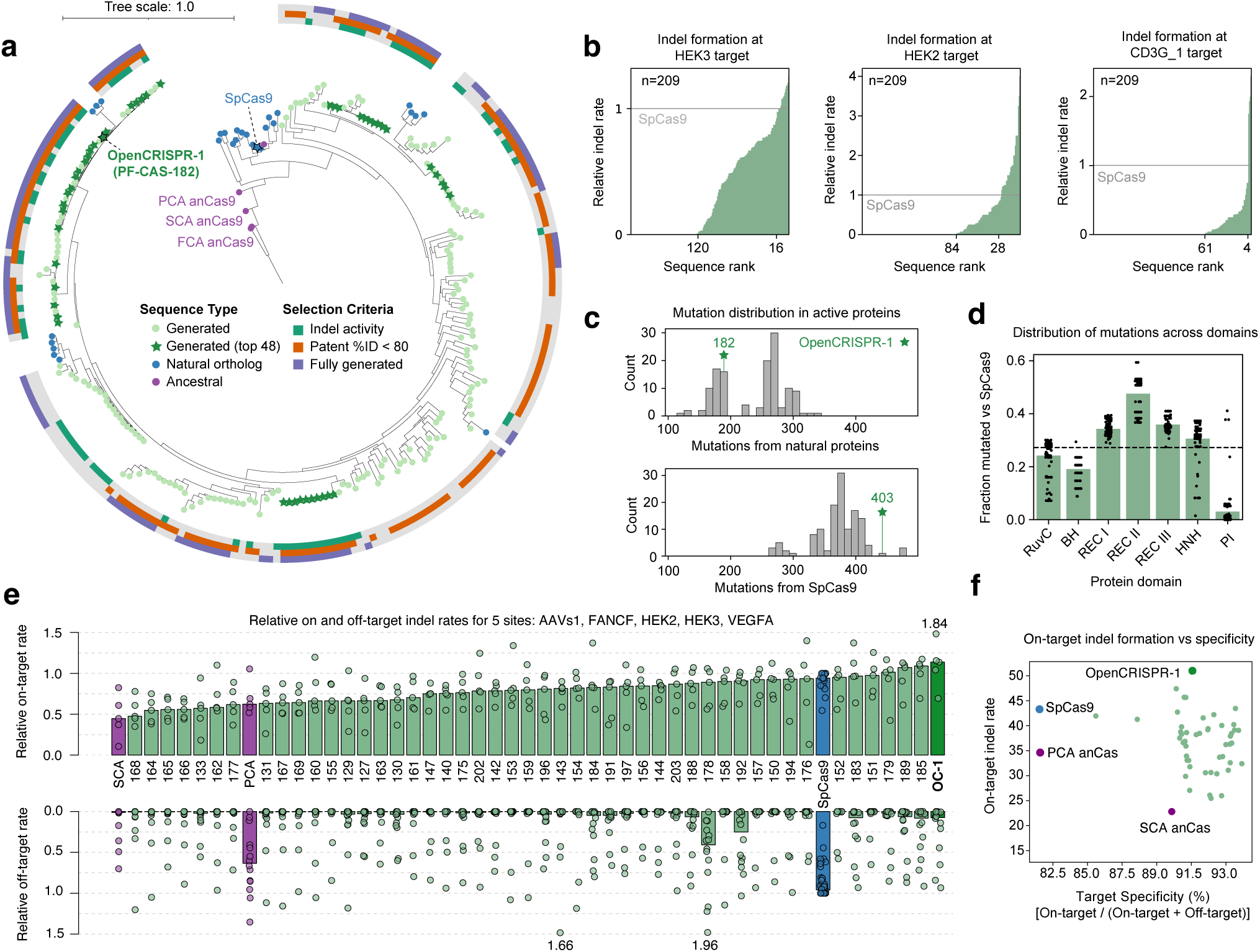
Generated nucleases function as gene editors in human cells. a) Phylogenetic tree of natural Cas9 proteins, ancestral reconstructions, and generated effector proteins near SpCas9. Annotations surrounding the tree indicate selection criteria used to identify 48 generated proteins for further characterization. b) Editing efficiency (indel rate relative to SpCas9) of 209 generated proteins across three target sites. Sequences are ordered according to relative indel rates, with the number of sequences showing activity and surpassing SpCas9 indicated on the x-axis. c) Mutational Levenshtein distances from the nearest natural protein in the CRISPR-Cas Atlas and SpCas9 for 131 generated proteins with observed editing activity. The Levenshtein distance is the minimal number of edits between two sequences, including substitutions, insertions and deletions. d) Domain-level mutational deviation from SpCas9 for 131 generated proteins with observed editing activity. The fraction mutated vs SpCas9 is measured as the Levenshtein distance between the two domains, normalized by the domain length for SpCas9. Horizontal line indicates the median whole-protein fraction mutated across all generated proteins. e) On- and off-target editing efficiency for natural Cas9s, ancestral reconstructions, and 48 generated proteins. Points correspond to on- or off-target editing at five sites (with three off-targets per site). Bars reflect the median of all on- and off-target editing. f) Comparison of on-target editing efficiency and specificity for natural Cas9s, ancestral reconstructions, and generated proteins. Specificity is calculated as the fraction of on-target editing over the sum of on- and off-target editing, tabulated separately for each off-target site then averaged. Points represent the average of on-target editing and specificity across the five targets. Generated proteins are able to achieve high on-target performance with increased specificity.

We next set out to explore whether our Cas9-like proteins were able to mediate genome editing in human cells. We human codon optimized and cloned the 209 sequences containing a C-terminal 1x SV40 NLS tag into CMV-driven expression plasmids. We initially tested for protein function using SpCas9 sgRNAs, which were predicted to be compatible with all Cas9-like proteins. We co-transfected HEK293T cells with the Cas9-like protein plasmid and sgRNA expression plasmids targeting one of three previously characterized target sites and inferred DNA repair outcomes after three days using a Sanger-sequencing based method (31). Across all three sites, we observed a wide range of editing efficiencies, with a subset of Cas9-like proteins showing activity on-par or higher than SpCas9 (Fig. 3b). In contrast with edit distance to SpCas9, language model scores were highly predictive of enzyme activity, separating active and inactive enzymes at the HEK3 target site with an AUC ROC value of 0.82 (Fig. S9). Among the set of active nucleases, we observed significant sequence deviation from SpCas9 and the nearest natural proteins within the CRISPR-Cas Atlas (Fig. 3c). Mutations were spread across most domains of the protein, while being enriched in Rec II and depleted from the PI domain (Fig. 3d). Cas9 proteins can tolerate large deletions in Rec II (32), which may be under less evolutionary constraint than other Cas9 domains.

For further characterization, we next selected 48 Cas9-like proteins that were fully generated (both N- and C-terminal domains), displayed moderate or high indel rates across one or more sites, and were substantially different from any natural or engineered enzyme in patent databases (lens.org) (Fig. 3a). We assayed the editing efficiency and specificity of these 48 proteins across a panel of previously characterized SpCas9 on- and off-target sites (n=5 and n=15, respectively) (33, 34), alongside SpCas9 and two ancestrally reconstructed variants that were recently shown to be active in human cells (35). Nuclease- and sgRNA-expressing plasmids were co-transfected in HEK293T cells, and DNA repair outcomes were measured after three days using next-generation sequencing of amplicons (NGS). We observed both high editing efficiency and specificity with many of our generated nucleases (Fig. 3e), with some even outperforming SpCas9 (Fig. 3f).

Our top hit, PF-CAS-182, displayed comparable levels of activity to SpCas9 at on-target sites (median indel rates of 55.7% vs 48.3%), while having a 95% reduction in editing at off-target sites (median indel rates of 0.32% vs 6.1%). Compared to the ancestral reconstructions, PF-CAS-182 was substantially more active (Fig. 3e), while having lower sequence similarity to SpCas9. PF-CAS-182 was 1,380 aa in length and considerably diverged from both SpCas9 (403 mutations) and any natural protein in the CRISPR-Cas Atlas (182 mutations). Alignment to NCBI-nr revealed top-hits to proteins from several Streptococcus spp. (S. cristatus, S. oralis, S. sanguinis), but none exceeding a sequence identity of 86.3%. PF-CAS-182 lacked previously identified immunodominant and subdominant SpCas9 T cell epitopes for HLA-A*02:01, suggesting it may be less immunogenic than SpCas9 (36) (Fig. S10). Based on its performance and novelty, we nominated PF-CAS-182 as the OpenCRISPR-1 protein.

Sequence alignment indicated significant sequence divergence between OpenCRISPR-1 and SpCas9, with a sequence identity of 71.7%. Template-based AlphaFold2 (29) predictions of the catalytic state (37) of OpenCRISPR-1 illustrated that the majority of the mutations were concentrated at the solvent exposed surface of the protein, with only a fraction located at the protein-nucleic acid interface (Fig. S11a-b). The majority of critical nucleic acid coordinating residues and nuclease site components were preserved, demonstrating the capability of the model to accurately constrain all necessary catalytic and interaction sites (Table S1). In addition to point mutations, OpenCRISPR-1 contained two loop insertions in the REC1 and HNH domains. We speculate that the nine-residue positively charged insertion in the REC1 domain may establish novel stabilizing phosphate-backbone interactions with both the repeat:anti-repeat segment of the guide RNA, as well as the PAM-proximal region of the target DNA (Fig. S11c). This insertion is analogous to sequence graftings of positively charged loops between natural Cas9 orthologs to boost activity (38). The four-residue insertion in the HNH domain is fully compatible with all the experimentally elucidated catalytic states (37) and could potentially have a stabilizing effect on the cleavage checkpoint state (Fig. S10d).

To more thoroughly characterize OpenCRISPR-1, we used NGS to screen 98 previously characterized SpCas9 target sites harboring either canonical or non-canonical PAMs (13, 39) (Table S3). After quality control, we were able to characterize nuclease performance at 92 of these sites (NGG PAMs, n=49; non-NGG PAMs n=43). In agreement with our prior experiment, OpenCRISPR-1 displayed comparable levels of on-target activity across sites bearing the NGG PAM (Fig. 4a-b) and resulted in a similar distribution of DNA repair outcomes (Fig. S12). Interestingly, OpenCRISPR-1 exhibited several-fold reduction in activity at genomic sites bearing a mismatch in the PAM (Two-sided Wilcoxon-rank-sum p-value=0.0005) (Fig. 4c). These results suggest that OpenCRISPR-1 has comparable activity to SpCas9 at on-target sites, while avoiding double-strand breaks at sites with mismatches in either the PAM or target regions.

**Figure 4.**
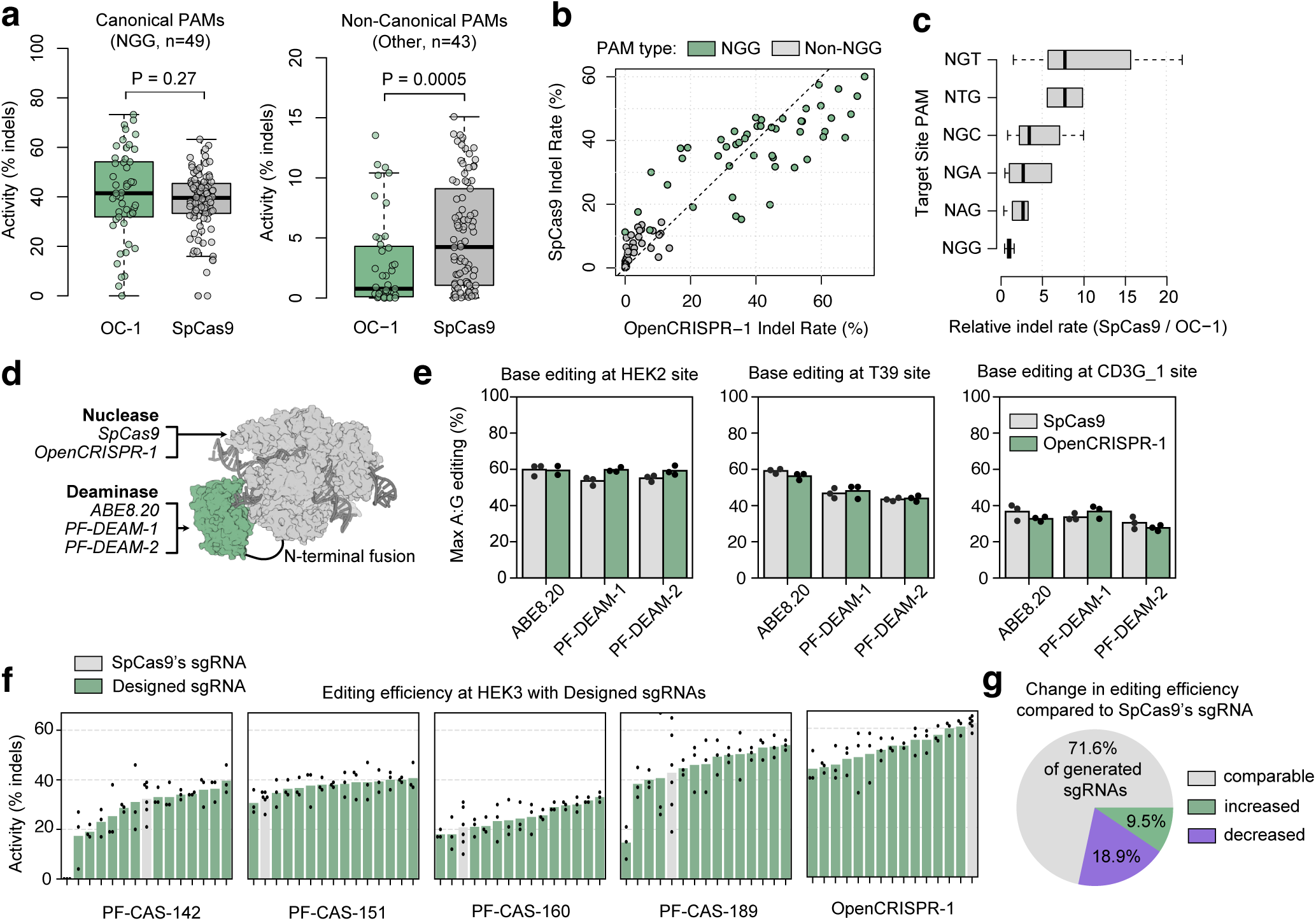
Characterization of OpenCRISPR-1 across PAMs, guides, and base editing. a-b) On-target editing efficiency (indel formation) of OpenCRISPR-1 protein at canonical (NGG, n=49) and non-canonical PAMs (n=43). OpenCRISPR-1 exhibits comparable activity at targets with an NGG PAM, but lower editing at sites lacking an NGG PAM. c) Relative activity of SpCas9 to OpenCRISPR-1 across sites with different PAMs (NGG, n=49; NGC, n=11; NGT, n=10; NGA, n=10; NAG, n=9; NTG, n=2; NCG, n=1). d) Adenine base editors were created by attaching deaminase domains to the N-terminus of OpenCRISPR-1 and SpCas9 nickase variants (D10A mutation for both proteins). e) Adenine base editing efficiency at three target sites. ABE8.20 is a highly active deaminase from directed evolution, while PF-DEAM-1 and PF-DEAM-2 were generated from language models. Across all target sites and with distinct deaminases, OpenCRISPR-1 nickase shows compatibility with base editing. f) Editing efficiency at HEK3 target site with designed sgRNAs (green) and SpCas9’s sgRNA (gray). Four of five generated proteins displayed increased editing efficiency with design sgRNAs. g) The majority of designed sgRNAs yield performance that is not significantly different from SpCas9’s guide, while a subset either significantly improves or worsens editing efficiency (T-test p-value < 0.05).

Next, we investigated whether OpenCRISPR-1 could be utilized in a base editing system. Base editors have emerged as powerful systems for modifying single nucleotides in the genome without the complications of generating double-strand breaks. We converted OpenCRISPR-1 to a putative target-strand nickase (containing D10A mutation) and fused it to a previously engineered adenosine deaminase (ABE8.20) commonly used in base editing (40) (Fig. 4d). We tested for editing in HEK293T cells using plasmid delivery with sgRNAs targeting three genomic loci containing adenines in the editing window (i.e. position 3-9 in the spacer). We observed robust A-to-G conversion with the OpenCRISPR-1 base editor on all three target sites (35-60% editing rate) which was comparable to an ABE8.20 base editor system using SpCas9 nickase (Fig. 4e) and without resulting in indel formation (Fig. S13).

Following our success, we next set out to engineer a fully synthetic base editor system, including the deaminase domain. Towards this goal, we trained models based on TadA-like proteins from UniProtKB (41) and BFD (29) and generated a series of synthetic adenine deaminases with 55-80% identity to any known natural sequence, including the E. coli TadA from which prior work has evolved potent editors (33, 40). Initial screening experiments with SpCas9 nickase revealed that a subset of generated deaminases were active at several targets in the human genome. We then tested two of our most active deaminases (PF-DEAM-1, PF-DEAM-2) fused to the N-terminus of either SpCas9 or OpenCRISPR-1 nickases. Notably, our generated deaminases showed A-to-G editing levels comparable to ABE8.20 with both nickase scaffolds, while producing minimal bystander edits (Fig. S14).

While all of the experiments up to this point had used the SpCas9 sgRNA scaffold, we reasoned that this might not be optimal for OpenCRISPR-1 and other generated Cas9-like proteins, which contained hundreds of mutations that could potentially disrupt RNA interactions. Using the gRNA model described previously, we designed fourteen generated sgRNAs for each of five generated Cas9-like proteins (including OpenCRISPR-1) and tested for editing in HEK293 cells at one target site (HEK3). Generated sgRNA sequences exhibited high sequence conservation compared to SpCas9’s sgRNA (Fig. S15), with most mutations occurring in flexible regions of the secondary structure (e.g. loops or linker regions). Overall, we observed enhanced editing with 42 designed sgRNAs relative to SpCas9’s sgRNA (Fig. 4f), including significant improvements for 7 sgRNAs for two of five variants (Fig. 4g). Designed sgRNAs were generally compatible with SpCas9 but did not yield any significant improvements in editing efficiency (Fig. S16d-f). Notably, we found that OpenCRISPR-1, which had shown consistently high editing efficiency in prior experiments, performed similarly with a designed sgRNA or SpCas9’s sgRNA. These results suggest that this generated protein may be applicable either as part of a fully generated gene editor or as a drop-in replacement for SpCas9 in existing editing systems, but further evaluation of the on- and off-target behavior of the fully generated construct is needed.

## Discussion

Gene editing technologies adapted from natural prokaryotic antiviral systems have enabled precise, programmable manipulation of genetic material across research, therapeutic, and industrial applications. While evolution has created a massive diversity of CRISPR-Cas proteins, identifying the best natural protein for a given application (if it exists) remains a major bottleneck in the design of more advanced gene editing systems. Generative language models for DNA (42) or proteins (18) offer an alternative paradigm, wherein models learn from natural diversity and can be steered towards the most promising regions of sequence space. This approach allows us to diversify existing lineages of interest or explore regions of sequence space that were never visited by evolution. In this work, we focused on generating Type II effectors in the phylogenetic neighborhood of SpCas9, ultimately yielding the OpenCRISPR-1 editing system. Our results suggest that OpenCRISPR-1 may provide a viable alternative to SpCas9 for use in gene editing technologies, with similar editing behavior and compatibility with systems like base editing. In the future, it will be important to examine OpenCRISPR-1 activity across a range of experimental conditions, cell types, and delivery methods in order to more thoroughly characterize robustness (9, 43, 44). Ultimately, we anticipate that optimization with language models offers the most promising path towards unlocking improvements in enzyme performance in these contexts.

With the aim of designing novel gene editing systems, we curated the CRISPR-Cas Atlas, which to the best of our knowledge represents the largest documented resource of CRISPR systems. Large, high-quality datasets like the CRISPR-Cas Atlas are critical for distilling the general learnings of pre-trained protein language models into a functional blueprint for design. Although we focused on Type II effector proteins in this work, early experiments suggest that effectors from other Class 2 systems (e.g., Cas12a, Cas12f, Cas13) may be amenable to the same approach. In some cases, these alternative systems have unique properties that would benefit gene editing applications (e.g., reduced size of Cas12f or RNA interference of Cas13). Aside from fine-tuning protein language models for generation, we envision that the CRISPR-Cas Atlas could be used to model specific properties of gene editors, such as nuclease size, PAM preference, tracrRNA compatibility, thermostability, or temperature-dependent activity. For instance, a model to predict PAM preference could enable efficient engineering of target- or allele-specific editors. The capability of generative language models to produce diverse, highly functional nuclease proteins, as demonstrated in this work, provides a foundation from which to pursue these fit-for-purpose editors.

Computational protein design has advanced considerably in recent years with the development of increasingly sophisticated deep learning algorithms. These improvements have largely been achieved through integration of more powerful tools into design pipelines that have remained unchanged for decades (45). Specifically, the design of protein function typically requires an explicit structural hypothesis that is translated into a set of constraints to guide a search for satisfying sequences (46). This approach has largely reduced some design problems, such as de novo design of protein mini-binders (47), to practice. However, for the design of complex functions as embodied by the gene editors in this work, structure-based approaches do not offer a straightforward solution. By contrast, language models provide an implicit means of modeling protein function (and thus structure) through sequence alone (16). By sampling unconditionally from fine-tuned language models, we were able to generate diverse sequences that recapitulate key functional domains through confident structure predictions, with as little as 40% identity to the training dataset. Detailed structural analysis revealed that OpenCRISPR-1 maintained all of the key functional residues involved in nuclease activity, despite the model never being explicitly tasked with doing so. For experimental characterization, we focused on generating nucleases that were compatible with SpCas9’s PAM and tracrRNA. Even under these constraints, we found that language models can generate nucleases that are hundreds of mutations away from any natural protein, yet function on par with SpCas9. In the future, it should be possible to relax these constraints and screen a broader diversity of nucleases in tandem with designed sgRNAs.

## Data availability

All data used to create the CRISPR-Cas Atlas are available in their original form via the IMG/M, ENA, and NCBI databases. Pre-trained protein language models (ProGen2 and ESM-2) used in this work are available on Github. The OpenCRISPR-1 protein and guide RNA sequences are provided at https://github.com/Profluent-AI/OpenCRISPR.

## Acknowledgements

We would like to thank Jakob Russel and Rafael Pinilla Redondo for assistance with the CRISPR-Cas discovery pipeline, Michael Marytn for help processing NGS data, and Dylan Morris for advice on project conception.

## Author contributions

A.M. and A.J.M. conceived of the project. A.M. supervised the project. S.N. curated data for the CRISPR-Cas Atlas. J.A.R., A.B., and J.B. performed machine learning model training and sequence generation. J.A.R. and A.B. selected generated proteins for experimental characterization. J.G. and P.C. designed wet lab experiments. J.G., R.H., J.R., J.Y., and E.H. performed wet lab characterization of Cas9-like proteins and base editors. M.P. performed structural analysis of OpenCRISPR-1. A.M., J.A.R., and S.N. wrote the first draft of the manuscript. J.A.R. and S.N. designed manuscript figures. All authors wrote and/or reviewed the final draft of the manuscript.

## Safety and ethics

Computational protein design carries the dual potential to accelerate development of novel therapeutics and other society-improving molecules, while providing parallel capabilities for nefarious uses, such as engineering of bioweapons. When bolstered by current and future iterations of generative AI, these capabilities are heightened and expected to grow further. The global protein design community has begun to establish appropriate regulations and guidelines towards the continued beneficial development and application of these technologies. In support of these efforts, J.A.R. and A.M. have joined as signatories on a set of community values, guiding principles, and commitments for the responsible development of AI for protein design (https://responsiblebiodesign.ai/). Gene synthesis represents a critical step in the actualization of designed protein sequences. The International Gene Synthesis Consortium (IGSC) unites major gene synthesis providers under a commitment to screen all incoming orders against known pathogens and potentially dangerous sequences. As a concrete step towards safe application of protein design technology, all gene synthesis work in support of the present study was performed with IGSC members. For all protein design projects, we urge researchers to maintain ethical oversight throughout project initiation, experimental characterization, and subsequent deployment phases to ensure safety and avoid unintended harmful outcomes.

By releasing OpenCRISPR-1, we aim to accelerate the development of one-and-done cures for genetic diseases. As with other genome editing technologies, OpenCRISPR-1 presents a set of dual-usage concerns, which have been discussed at length within the genome editing community. Through the characterization presented here, we have found no evidence that OpenCRISPR-1 presents risks exceeding those posed by SpCas9 or other naturally derived effectors. We are firmly committed to safe and ethical usage of our technology, and will continue to monitor and engage in global scientific discussions regarding the appropriate application of OpenCRISPR-1 and future editors. However, we believe that the potential benefits of accessible, effective gene editing for therapeutic, agricultural, and research applications outweigh these concerns.

## Methods

### Data curation

#### Curating an atlas of CRISPR-Cas operons

We downloaded 26.2 Tbp of assembled microbial genomes and metagenomes from diverse public data sources including IMG/M (48), ENA (49), NCBI (50), and assembled metagenomes from the human microbiome (51). To reduce data from highly over-represented taxa in NCBI we selected up to 200 assemblies per species and up to 2000 per genus. This resulted in a final dataset of 134,182 assembled metagenomes and 509,320 assembled microbial genomes and metagenome-assembled genomes (MAGs). To efficiently mine this data, we used a combination of CRT (52) and PILER-CR (53) to rapidly identify contigs >5 kbp containing a CRISPR array and extracted up to 20 kbp of genomic context flanking each CRISPR. Prodigal-gv v2.11.0 (54) (options: -p meta) was used to identify protein coding genes. Cas proteins were annotated using hmmsearch v3.4 (55) with a custom database of non-redundant profile HMMs from the CRISPRCasTyper (56) combined with an expanded set of Cas9 and Cas12 HMMs constructed for the current manuscript (n=480). Adjacent proteins were clustered into operons having at least one CRISPR array, at least one Cas protein, and maximum of 3 consecutive unannotated genes. A total of 1,246,163 putative CRISPR-Cas operons were identified with mean length of 7,701 bp and containing on average 5.71 total genes and 4.95 annotated Cas genes. We refer to this resource as the CRISPR-Cas Atlas.

#### Diverse CRISPR-associated families

For modeling diverse CRISPR-associated protein families, we utilized a curated subset of 5,100,022 full-length proteins (n=1,123,633 unique) that confidently matched a Cas protein profile HMM (bitscore >= 40, E-value < 1e-5, HMM alignment coverage >70%) and were not truncated by a contig edge. For modeling Cas9 proteins, we utilized a subset of 238,917 Cas9-annotated proteins (n=64,739 unique) from the CRISPR-Cas Atlas. Proteins were clustered with MMseqs2 v15.6f452 (57) at 30%, 50%, 70%, and 90% amino acid identity over 80% of the length of both query and target sequences. MMseq2 cluster assignments were highly consistent with the HMM-annotated protein family labels (99.72% agreement rate within 30% identity clusters having >1 member).

#### Identification of tracrRNAs

tracrRNAs were identified using a database of 1,747 covariance models that broadly cover Cas9 tracrRNA diversity (58). Covariance models were searched against Type II CRISPR operons using cmsearch from the infernal v1.1.5 package (59) (options: –notrunc –cpu 1 -g). To improve pipeline efficiency and accuracy, we masked protein coding regions with Ns prior to running cmsearch. We retained alignments with an E-value < 1e-3, resulting in predictions for 258,036 Type II CRISPR-Cas operons. Anti-repeats were identified by aligning the direct repeat from the CRISPR array to the predicted tracrRNA using blastn v2.15.0+ (options: -word_size 5 -dust no) (60). We retained putative anti-repeats with an alignment length of at least 15 nt which started within 5 bp of the start of the tracrRNA. Terminators were identified based on the presence of a TTTT motif within the last 14 nucleotides of the predicted tracrRNA sequence. tracrRNAs for 117,802 CRISPR-Cas operons were classified as high-confidence based on having: (1) a predicted anti-repeat, (2) a terminator, (3) a cmsearch E-value < 1e-5, and (4) not overlapping a predicted protein-coding region. High-confidence tracrRNA predictions were used as input to train the gRNA model.

#### Identification of PAMs

PAMs were inferred for the CRISPR-Cas operons with PAMpredict v1.0.2 (61) using a database 15,663,652 viral sequences from IMG/VR v4 (62) and 699,973 plasmid sequences from IMG/PR (63). To improve sensitivity, we pooled together CRISPR spacers from (near)identical Cas proteins clustered using MMseqs2. Within each Cas protein cluster, CRISPR repeats and spacers were clustered using CD-HIT (options: cd-hit-est -c 0.95 -s 1.0) (64) in order to consistently orient arrays from different CRISPR operons (which may be found on the top or bottom strand) and to remove duplicate spacers.

### Modeling of CRISPR-Cas sequences

#### CRISPR-associated protein language model

To model CRISPR-associated proteins, we fine-tuned language models on the full set of proteins from the CRISPR-Cas Atlas. This dataset was clustered at 50% sequence identity to create train, validation, and test splits with 1,077,601, 22,985, and 23,047 sequences, respectively. Particular families were over-represented in the dataset (e.g., Cas1 and Cas2) and had the potential to skew training if sequences were sampled uniformly. To account for these distributional biases, we sampled a family-balanced dataset for each training epoch. Additionally, to avoid over-sampling highly similar sequences, we balanced sampling within families over 90%-identity clusters according to the inverse-log of their size. Sequences were provided to the model in forward (N-to-C) and reverse (C-to-N) directions. We initiated fine-tuning from the pre-trained ProGen2-base model, which was further trained for a maximum of five epochs with an effective batch size of 8,200 residues. Learning was increased linearly over 70,000 warmup steps to a maximum value of 5e-5, then decayed according to an inverse-square-root schedule. The best model checkpoint was selected according to validation loss.

We generated four million sequences using the CRISPR-Cas PLM. Two million were sampled directly from the model, reflecting the learned distribution over CRISPR-Cas families. Then, to guide the model towards generation of sequences belonging to the 45 most highly represented families, we generated an additional two million sequences (45,000 per family) by initiating generation with a prompt from a natural protein. Prompts were constructed by randomly selecting a natural protein from a family-of-interest then uniformly sampling 1-50 residues from either the N- or C-terminus. Sequence generation direction was balanced evenly between N-to-C and C-to-N. All sequences were generated with temperature *T* = 0.5 and nucleus probability *P* = 0.95.

After generating the full set of four million sequences, a series of filters were applied to ensure only realistic proteins were used to characterize the model’s generative capabilities. Any sequences containing non-canonical or unknown amino acids were discarded. Degenerate sequences were filtered according to k-mer repetition, such that no amino acid motif of 6/4/3/2 residues was repeated 2/3/6/8 times consecutively, respectively. These criteria were satisfied by over 99.5% of natural proteins from the CRISPR-Cas Atlas yet removed dominant failure modes among generated sequences. Additionally, all sequences were filtered according to coverage within CRISPR-Cas families. Using BLAST, all generated sequences were aligned against 90% identity cluster representatives from the CRISPR-Cas Atlas. Sequences were retained if the top-scoring alignment (excluding exact matches) had greater than 80% query and target coverage, as well as a BLAST score greater than 50. Additionally, all sequences were aligned to a database of CRISPR-Cas profile HMMs using HMMER (55). Sequences were retained if the best-matching profile HMM had query and target coverage greater than 80% and a score greater than 50. Any sequence not satisfying either the BLAST or HMMER criteria were discarded. After applying all of the filtering criteria, two million generated sequences remained.

#### Cas9 protein language model

To more explicitly guide generation towards Type II effector proteins, we fine-tuned another language model on the subset of Cas9 proteins from the CRISPR-Cas Atlas. This dataset was clustered at 90% sequence identity to create train and validation splits with 58,553 and 6,186 sequences, respectively. We performed the same fine-tuning strategy as described for the CRISPR-Cas PLM using this dataset, sampling according to the inverse-log of 90% cluster sizes. We generated one million sequences directly from the Cas9 PLM and 200,000 using the prompting strategy described previously. All sequences were generated with temperature *T* = 0.5 and nucleus probability *P* = 0.95, then filtered according to the amino acid composition, k-mer repetition, and coverage criteria described previously.

#### Analysis of generated Cas9-like proteins

We calculated the percent identity and edit distance of all generated Cas proteins versus SpCas9 (UniProt ID: Q99ZW2), natural Cas9 proteins from the CRISPR-Cas Atlas (n=238,917), and proteins listed in patent claims from the Lens.org database (release date=2024-02-01, n=7,518,295). Alignments were performed using diamond blastp (version: v2.1.9, options: –very-sensitive) (65) and percent identity was calculated over the full length of the query protein sequence. To identify protein domains, Pfam HMMs were downloaded from InterPro (release 2023-09-12) (66) and aligned to Cas9 proteins using hmmsearch (options: –cut_tc).

A phylogeny was constructed for generated Cas9 proteins (n=1,000,000) combined with natural from the CRISPR-Cas Atlas (n=238,917) and UniRef100 (n=9,564), biochemically characterized Cas9s from Gasiunas et al. (n=79) (5), and a set of Cas9s used in genome editing (n=15) (25). All proteins were clustered at 40% sequence identity using MMseqs2 (n=15,465). We constructed a multiple sequence alignment of cluster representatives using FAMSA v2.2.2 (67) that was trimmed using trimal v1.4.1 (options: -gappyout) (68) and pruned by removing 1% of sequences with the lowest alignment coverage. The final tree was constructed for remaining sequences (n=15,340) using FastTree v2.1.11 (69) and visualized using iTOL (70).

#### Guide RNA design model

We formulated the design of guide RNAs for a given type II effector protein as a sequence-to-sequence generation task. From the CRISPR-Cas Atlas, we collected the subset of Type II systems with identified tracrRNA and crRNA components. This subset was split according to 90% protein sequence identity clustering to form training and validation datasets with 99,822 and 12,390 data points, respectively. We constructed training examples by concatenating the tracrRNA and crRNA in a random order with delimiters indicating the boundaries of each. We then trained an encoder-decoder model to generate the concatenated gRNA sequences from their associated effector protein. For the encoder, we used a pre-trained ESM-2 model (8M parameters) with frozen weights, followed by a single bidirectional transformer layer. The RNA sequence was then decoded by three autoregressive transformer decoder layers with causal self-attention and cross-attention to the encoder embeddings. In total, the model had 700K trainable parameters. The gRNA model was trained for 40 epochs with an effective batch size of 64. The learning rate was increased linearly over 4,000 warmup steps to a maximum value of 1e-4, then decayed according to an inverse-square-root schedule.

To evaluate the capabilities of the gRNA design model, we generated a set of sgRNA sequences for ten Type II effector proteins that have previously been used as genome editors (SpCas9, ScCas9, St1Cas9, St3Cas9, PrCas9, SaCas9, GeoCas9, Nme1Cas9, Nme2Cas9, and CjCas9). For each protein, we generated 128 pairs of crRNA repeat and tracrRNA sequences with sampling temperature *T* = 1.0. The model occasionally produced truncated or non-terminating RNA sequences, which was addressed by discarding all sequences with concatenated length above and below the 90th and 10th percentile lengths of the set, respectively. We created single guide RNAs (sgRNAs) as previously described (5). We first aligned the generated crRNA repeat and tracrRNA sequences using blastn (options: -word size 4 -dust no). The end of the repeat and beginning of the tracrRNA were then trimmed and joined together with a flexible GAAA linker, resulting in a 12-bp stem (formed by the repeat:anti-repeat) and a 4-bp tetraloop. Length outliers for the set of sgRNAs above and below the 90th and 10th percentiles, respectively, were again removed. To select a diverse set of sgRNAs, we clustered the sequences at 95% identity using MMseqs2, then selected the best-scoring representative from each cluster according to the model log likelihood until ten sequences were sampled. If less than ten clusters were found, the remaining sequences were selected solely based on the model score.

### Generation of gene editors

#### Constrained generation of proteins

To generate novel Type II effector proteins for experimental characterization, we employed a constrained generation strategy wherein we initiated generation by providing either the PAM-interacting domain (PID) or the N-terminal non-PID domains of SpCas9 to the model. By doing so, we reasoned that the generated sequences would maintain compatibility with the PAM and sgRNA preferences of SpCas9, facilitating direct comparison of protein activity in a common experimental setting. For PID-conditioned generation of N-terminal domain sequences, we decoded the sequence from C- to N-terminus, while for generation of novel PIDs we generated from N-to-C. In total, we generated 150,000 N-terminal domain sequences and 200,000 PID domains. All sequences were generated using a combination of sampling temperatures (*T* ∈ {0.5, 0.7, 1.0}) and nucleus sampling probabilities (*P* ∈ {0.5, 0.95}).

#### Selection of proteins for characterization

We began the process of selecting Type II effector proteins for experimental characterization by filtering out sequences we expected to be invalid. We first applied the amino acid composition and k-mer filters described previously to remove any degenerate sequences. Next, we required that the Cas9 PLM perplexity of any generated sequence was no more than 0.15 higher than SpCas9’s perplexity of 0.23. To enforce interoperability, we trained supervised models – applying a 2-layer MLP to the [CLS] token of the 650M-parameter ESM-2 model – that predicted whether a given effector protein’s tracrRNA and PAM matched that of SpCas9. We discarded all generated sequences that were predicted to be incompatible. Finally, we threaded generated PID sequences onto SpCas9’s structure and discarded any sequences that had mutations within 8Å of the DNA, to ensure that any potential PAM-recognizing residues were conserved.

We then combined all remaining generated N-terminal segments with both SpCas9’s PID and all remaining generated PIDs. We selected a diverse set of proteins that maximized a weighted average of ProGen2-base, ProGen2-xlarge, and Cas9 PLM log likelihoods. We also required that a subset of these proteins had at most 80% identity to any sequence in the CRISPR-Cas Atlas and/or the Lens Patent Database.

#### Design of sgRNAs

For a subset of generated Cas9-like proteins, we designed sgRNAs using the procedure described previously for natural genome editing proteins. We generated 400 pairs of tracrRNA and crRNA sequences, from which 15 sgRNAs were selected for experimental testing according to the same scoring and clustering criteria.

#### Design of deaminases for base editing

Deaminase proteins for base editing were designed according to a similar methodology as described for Cas9-like proteins. We curated a dataset of TadA-like proteins by searching UniProtKB (41) and the AlphaFoldDB (71) for proteins with enzyme commission (EC) number 3.5.4.33 (denoting tRNA adenine-34 deaminase function), yielding 34,575 sequences with 28,664 having predicted structures. These sequences were clustered at 50% sequence identity and the representatives were used as seeds to search for additional unannotated TadA-like proteins in the UniProt, BFD, and MGnify databases using MMseqs2. In total, a set of 77,632 TadA-like sequences were collected and used to fine-tune the ProGen2-base language model. Using the predicted structures, we additionally fine-tuned the ProteinMPNN (14) structure-conditioned sequence design model. We then generated two million unique TadA-like sequences, which were filtered according to the amino acid composition, k-mer repetition, and sequence complexity filters described previously, yielding 1.2 million viable deaminases. These were further clustered at 80% sequence identity, and the best-scoring sequence from each cluster according to the fine-tuned language model was retained. Of the remaining 263,000 sequences, we stratified selection according to identity to natural proteins and selected the best-scoring 3,500 sequences from subsets with <60%, 60-70%, and 70-80% identity to natural proteins (10,500 total). Using ESMFold (17), we predicted structures for the remaining sequences and discarded any with pLDDT and PAE below 86 and 0.88, respectively, yielding 9,100sequences. From this set, we selected 62 proteins for experimental characterization, balancing for language model scores (pre-trained ProGen2, fine-tuned ProGen2, and fine-tuned ProteinMPNN), novelty (identity to natural proteins), and diversity amongst the selected sequences. From this set, we identified two highly active proteins, which were further optimized through redesign of non-active site residues using proseLM (manuscript in preparation), yielding the deaminases referred to as PF-DEAM-1 and PF-DEAM-2.

### Characterization of gene editors

#### DNA oligonucleotides and plasmid assembly

Oligonucleotides used in this study were synthesized by IDT with standard desalting. All natural and AI-generated nuclease and deaminase sequences were purchased as synthetic gene fragments (Twist Bioscience), human codon-optimized using the Twist codon optimization tool, and cloned into CMV-driven expression plasmids using HiFi DNA Assembly (New England Biolabs). Single-guide RNA (sgRNA) sequences were cloned into a human U6 (hU6)-driven expression plasmid that also contains a CMV-driven GFP transfection reporter using HiFi DNA Assembly (New England Biolabs). All plasmids were sequence-verified by whole plasmid Nanopore sequencing (Primordium) prior to downstream applications.

#### HEK293T cell culture and transient transfection

HEK293T cells (ATCC) were cultured at 37°C and 5% (v/v) CO2 in high glucose DMEM with 4 mM L-glutamine, 1 mM sodium pyruvate and phenol red pH indicator (Gibco), supplemented with 10% FBS and 1X penicillin-streptomycin. 24 hours prior to transfection, cells were seeded at a density of 1x103 cells/well in 96-well tissue culture-treated plates (Nunc™ Edge™, Thermo Fisher Scientific).

For each transfection well, 50 ng of sgRNA plasmid and 50 ng of nuclease or base editor-expressing plasmid were added to 5 *µ*L of Opti-MEM (Gibco). 0.2 *µ*L of TransIT®-2020 transfection reagent (Mirus Bio) was diluted into 4 *µ*L of Opti-MEM. Plasmid and TransIT®-2020 mixtures were combined, incubated for 15-30 min at room temperature, and added to HEK293T cells in a dropwise manner. Plates were gently rocked to mix and incubated for 72 hours.

#### Target site selection

Cas9 on- and off-target sites were selected based on previous publications and are listed in Table S2 (72). The expanded set of 98 sites (NGG PAMs: n=52; NAG PAM:, n=10, Other PAMs: n=36) was selected from previous publications and is listed in Table S3 (39).

#### Targeted-amplicon sequencing and analysis

Cell lysates were generated by washing the cells with 1X PBS and adding 25 *µ*L of lysis buffer (100 mM Tris-HCl, pH 7.5; 0.05% SDS; 25 *µ*g/mL Proteinase K) per well. Plates with lysis buffer were incubated at 37°C for 1 hour, and then 25 *µ*L of nuclease-free water was added per well. The lysates were transferred to 96-well PCR plates and boiled at 98°C for 15 min.

For short read Illumina NGS analysis, target sites were amplified for indel quantification using a two-step PCR reaction. First, 5 *µ*L of lysate (corresponding to 1x103 cells) was used as a template for PCR (Invitrogen™ Platinum™ SuperFi II PCR Master Mix) with unique primer pairs containing an internal locus-specific region and an outer Illumina-compatible adapter sequence Tables S2 and S3. The resulting product was then diluted 1:100 in nuclease-free water and used as a template in a second PCR reaction targeting the outer-adapter sequence, appending unique indices (xGen UDI 10nt Primer Plates 1-16, IDT) to each amplicon for pooled sequencing. Amplicons were pooled 1:1 and sequenced on a NovaseqX with 2 x 151 paired end reads (Seqmatic). All amplicons across experiments included reference samples that were not treated with active nuclease to control for any variant reads relative to reference genome that are not due to gene editing.

After de-multiplexing, we obtained 25,004 to 5,711,684 reads per amplicon. Reads were trimmed and filtered using fastp (options: -unqualified_percent_limit 20 –average_qual 25 –length_required 120) removing 2-5% of reads per sample (73). Next, we ran CRISPResso2 v2.2.24 in “CRISPRessoBatch” mode with default parameters using the processed reads, amplicon sequences, and spacer sequences as input (74). For off-target sites, spacer sequences were edited to match the corresponding region on the amplicon. The indel rate was computed as the percentage of aligned reads having an insertion or deletion within one bp of the cleavage site. We excluded six amplicons which displayed low read quality, low alignment rates, high rates of substitutions, or high rates of editing at negative controls.

For Sanger sequencing analysis, locus-specific primers (Table S4) were used to amplify regions of interest from cell lysates by PCR (Q5® High-Fidelity DNA Polymerase). After PCR, the resulting amplicons were purified (Mag-Bind® RxnPure Plus, Omega Bio-tek), DNA yields were quantified (QuantiFluor® kit, Promega), and DNA concentrations were normalized to 2 ng/100 bp of amplicon length and submitted for Sanger sequencing with the appropriate forward PCR primer. We quantified the activity of base editors and nucleases using the software tools BEAT (75) and Synthego ICE v1.2.0 (31), respectively, with default parameters.

## Supplementary information

**Table S1.**
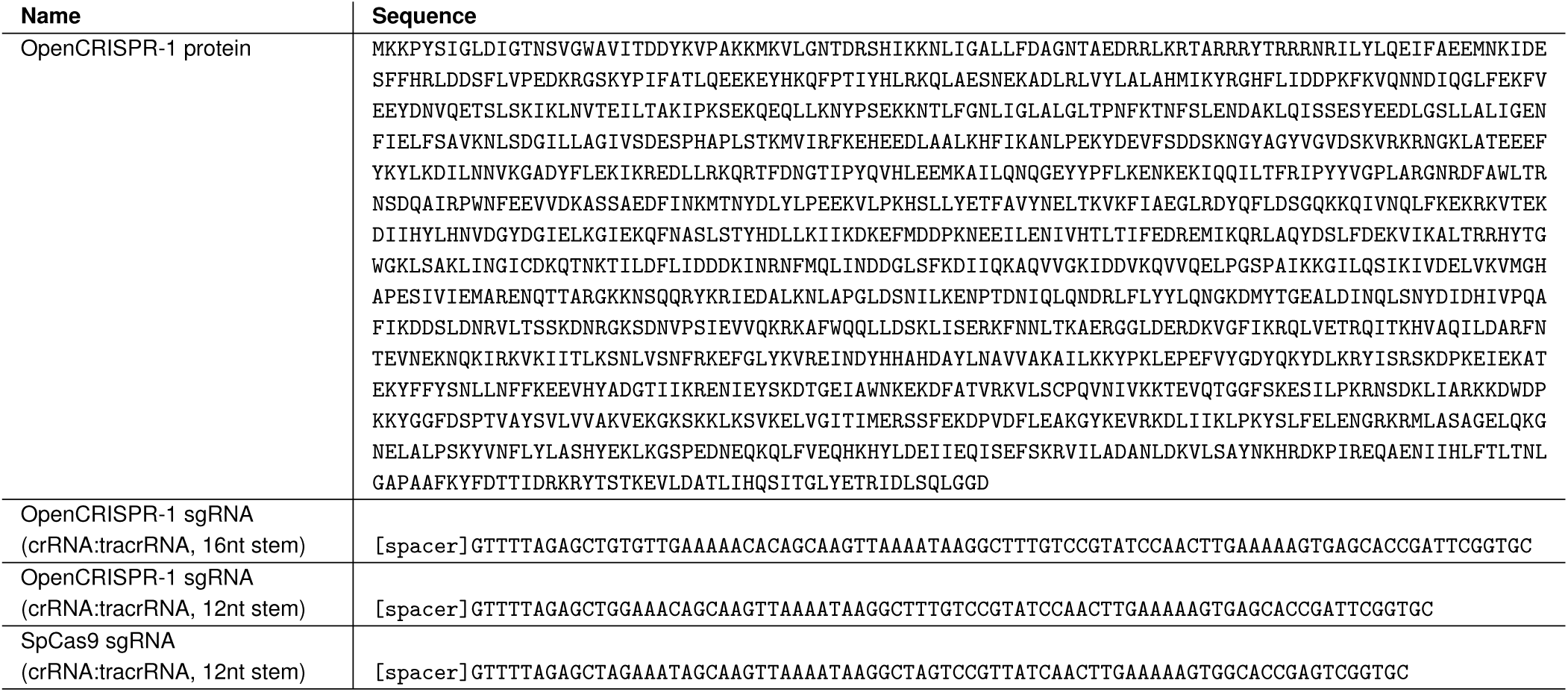
Sequences for OpenCRISPR-1 editing system.

**Table S2.** Guide sequences, target sequences and NGS primers for on- and off-target analysis.

**Table S3.** Guide sequences and NGS primers for expanded on-target characterization.

**Table S4.** Guide and primer sequences for Sanger sequencing analysis.

**Table S5.** Conservation of catalytic residues in OpenCRISPR-1.

**Fig. S1.**
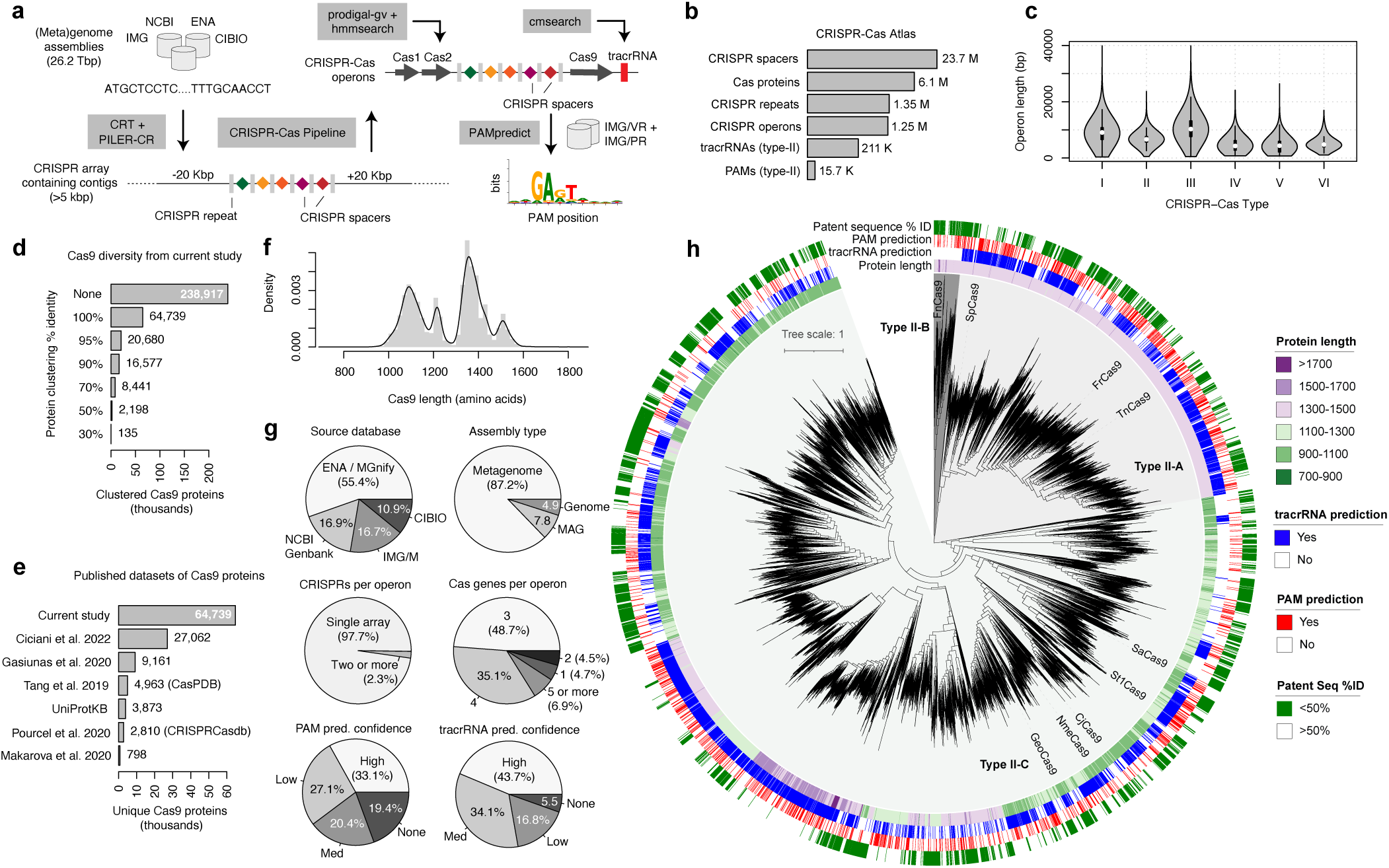
Formation of the CRISPR-Cas Atlas. a) Pipeline for discovery and annotation of 1.25M CRISPR-Cas operons from 26.2 Tbp of genome and metagenome assemblies. b) Summary of different entities across the CRISPR-Cas atlas. c) Distribution of operon lengths across CRISPR-Cas types. d) 238,917 Cas9 proteins were identified from Type II CRISPR-Cas operons and clustered using MMseqs2. e) Comparison of the number of unique Cas9 proteins compared to previously published datasets (5, 22, 61, 76–78). UniProtKB was queried in March 2024 using search term: gene = Cas9. f) Length of 64,739 unique Cas9 proteins from the CRISPR-Cas Atlas. g) Summary statistics across 64,739 CRISPR-Cas operons. h) Phylogenetic tree of 8,441 Cas9s clustered at 70% identity. Phylogenetic tree built using FastTree2 (69) and visualized using iTOL (70).

**Fig. S2.**
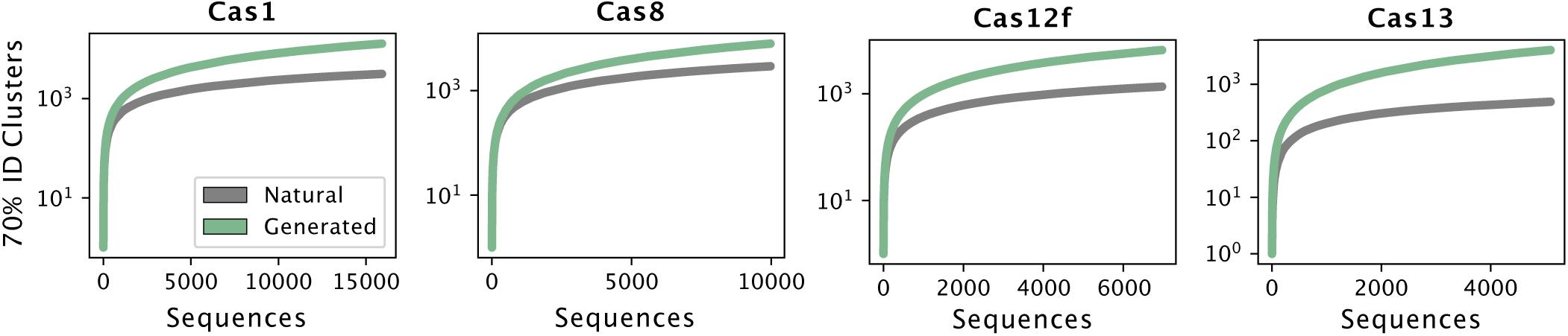
Protein family accumulation curves for natural and generated CRISPR-associated proteins. Cumulative cluster counts for four CRISPR-Cas families, showing the number of unique 70% identity clusters observed as increasing numbers of natural and generated sequences are sampled. Generated proteins access significantly greater diversity per sequence sampled, without showing saturation. Vertical axis is shown on a log10 scale. Each curve is the average of twenty independent rounds of sequence sampling.

**Fig. S3.**
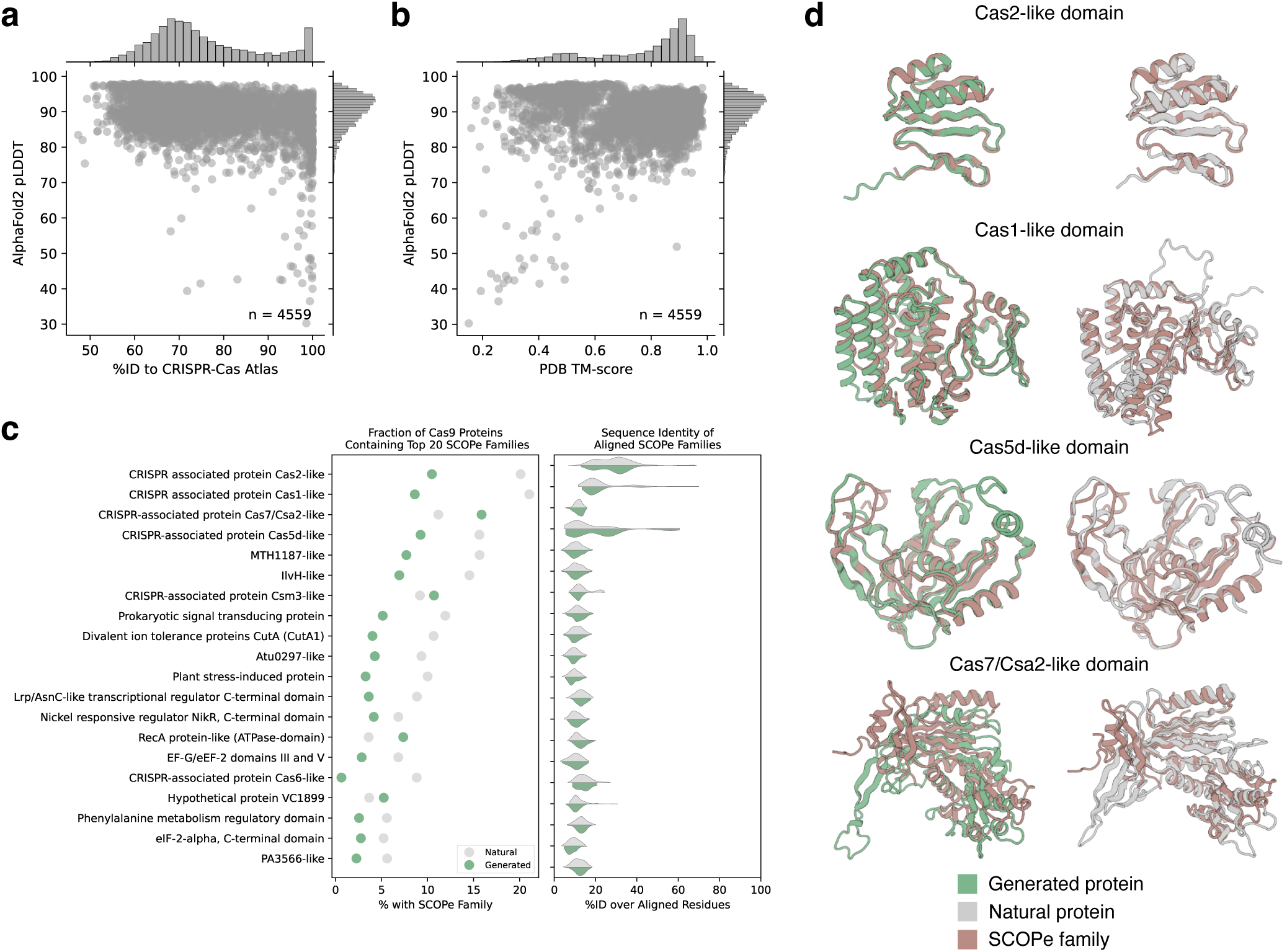
Structural composition of generated CRISPR-Cas proteins. Generated and natural CRISPR-Cas proteins were clustered at 70% identity using MMseqs2 (57). For both generated and natural proteins, random representatives of the largest 5,000 clusters were selected for structural analysis. Structures were predicted using the ColabFold (79) implementation (v1.5.2) of AlphaFold2 (29) using multiple sequence alignments (MSAs) from the ColabFoldDB (no templates). a) Generated sequences yield high confidence AlphaFold2 structure predictions despite significant sequence divergence from natural proteins. b) Predicted structures for generated proteins align well to experimentally determined structures from the PDB. c) Using Foldseek (80) (v8.ef4e960), predicted structures for generated and natural proteins were searched against the SCOPe database (30) (v2.08). Points in the left plot represent the fraction of generated (green) and natural (gray) proteins containing the twenty most commonly observed SCOPe families. Distributions in the right plot show the sequence identity over aligned residues between the generated and natural proteins and the best-matching SCOPe family structure. Overall, generated proteins were composed of similar structural components as compared to natural proteins. d) Examples of four most frequently observed SCOPe families (brown) aligned to generated (green) and natural (gray) proteins.

**Fig. S4.**
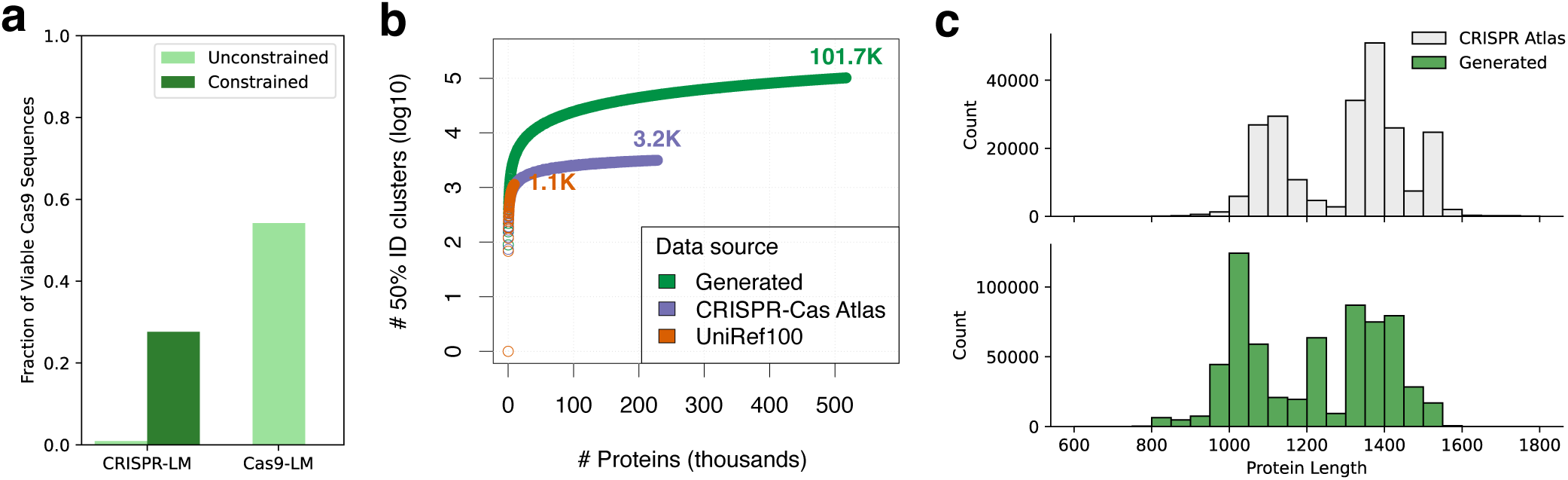
Cas9-like proteins generated by the CRISPR or Cas9 PLM. a) Fraction of viable sequences generated from each model, with or without a prompt. b) Accumulation curve of natural Cas9 and generated protein clusters. Vertical axis is shown on a log10 scale. Proteins are colored by their source (Generated: generated proteins from this study; CRISPR-atlas: Cas9 proteins recovered from CRISPR-Cas metagenome mining; UniRef100: Cas9 proteins identified from the UniRef100 database). c) Length distribution of generated proteins compared to naturals from the CRISPR-Cas Atlas.

**Fig. S5.**
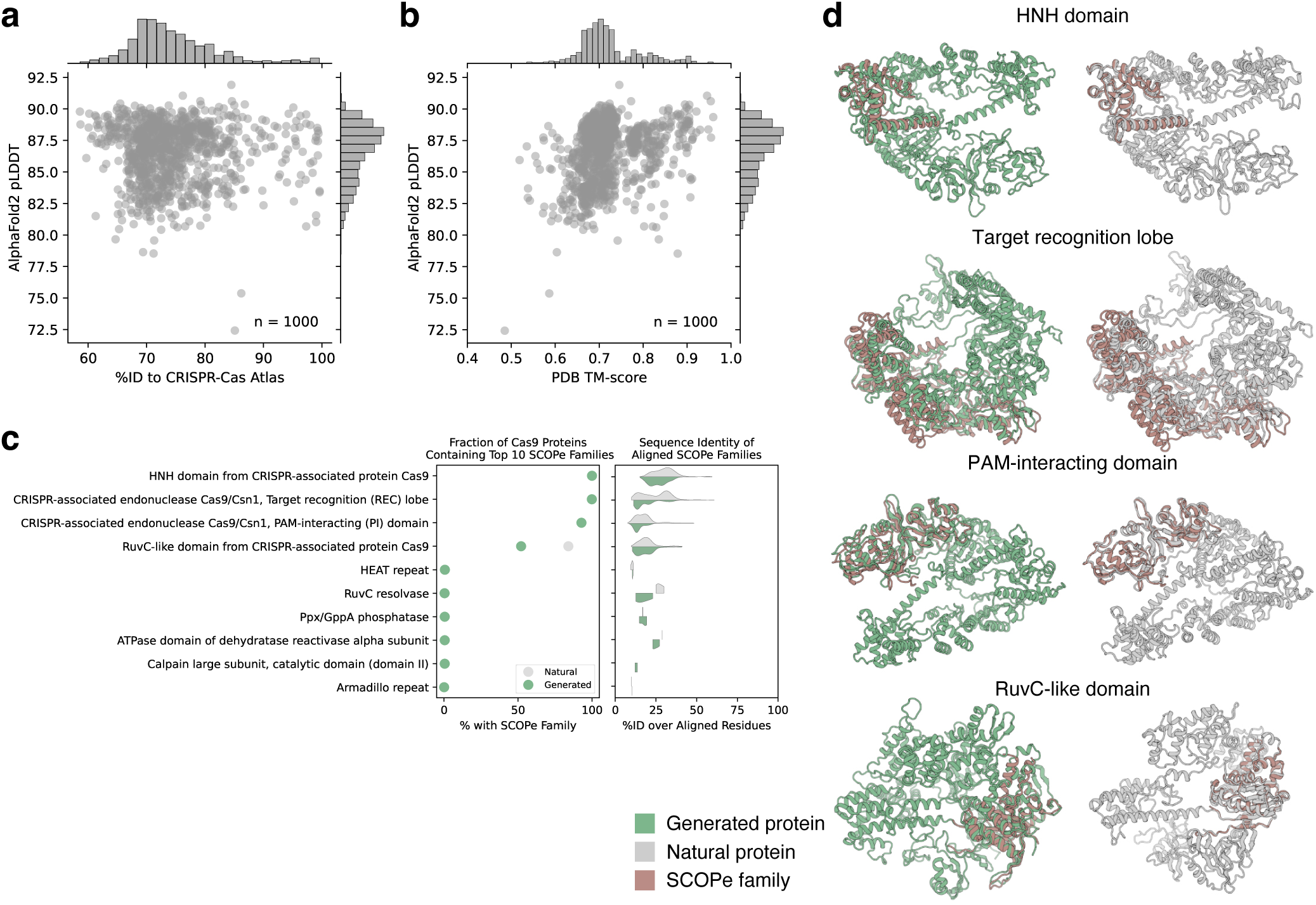
Structural composition of generated Cas9 proteins. Generated and natural Cas9 proteins were clustered at 70% identity using MMseqs2 (57). For both generated and natural proteins, random representatives of the largest 1,000 clusters were selected for structural analysis. Structures were predicted using the ColabFold (79) implementation (v1.5.2) of AlphaFold2 (29) using multiple sequence alignments (MSAs) from the ColabFoldDB (no templates). a) Generated sequences yield high confidence AlphaFold2 structure predictions despite significant sequence divergence from natural proteins. b) Predicted structures for generated proteins align well to experimentally determined structures from the PDB. c) Using Foldseek (80) (v8.ef4e960), predicted structures for generated and natural proteins were searched against the SCOPe database (30) (v2.08). Points in the left plot represent the fraction of generated (green) and natural (gray) proteins containing the ten most commonly observed SCOPe families. Distributions in the right plot show the sequence identity over aligned residues between the generated and natural proteins and the best-matching SCOPe family structure. Overall, generated proteins were composed of similar structural components as compared to natural proteins. d) Examples of four most frequently observed SCOPe families (brown) aligned to generated (green) and natural (gray) proteins.

**Fig. S6.**
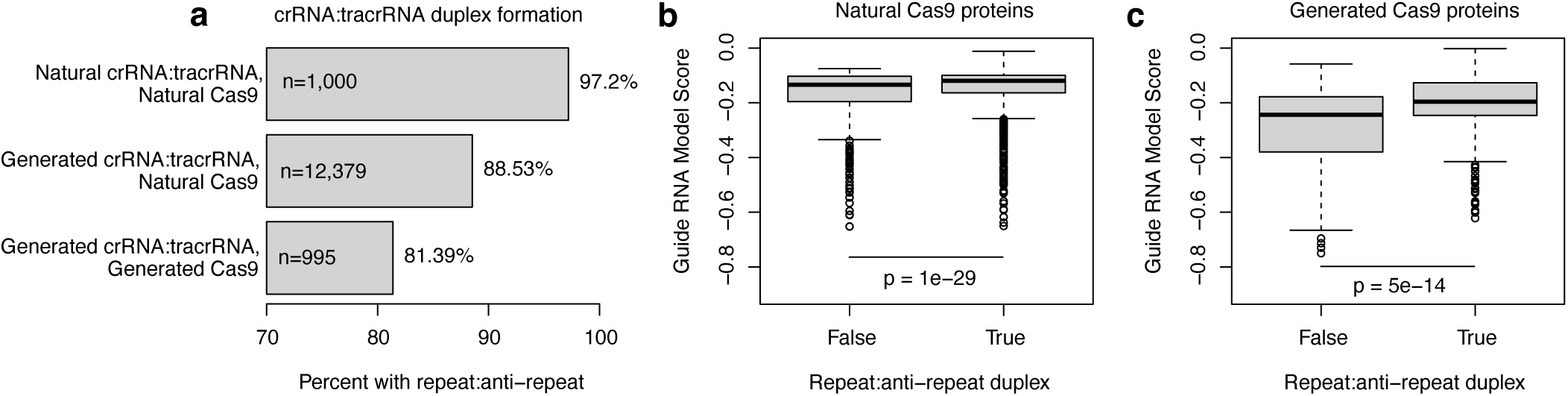
Duplex formation between natural and generated crRNA:tracrRNA pairs. RNAs were evaluated for three sets of sequences: natural RNAs for natural Cas9 proteins, generated RNAs for Cas9 natural proteins, and generated RNAs for generated Cas9 proteins. RNAs were aligned using blastn (-word_size 4, -dust no) and an anti-repeat was deemed present if 3’ end of crRNA (+ strand) aligned to 5’ end of tracrRNA (- strand). a) A high fraction of sequences from all three groups form anti-repeats. Anti-repeats %: Natural RNAs for natural proteins > Generated RNAs for natural proteins > Generated RNAs for generated proteins. b-c) The model score is somewhat predictive of whether a crRNA:tracrRNA will form a repeat for either RNAs generated for natural proteins, or for generated proteins. The model score alone is insufficient to accurately discriminate the classes (AUC values of 0.57 and 0.68, respectively), but the distributions are significantly different between classes (Wilcoxon-rank-sum test model P-values of 1e-29 and 5e-14, respectively). We conclude that the RNA models perform well, and can be used to design guide RNA scaffolds for either natural or synthetic Cas9 proteins.

**Fig. S7.**
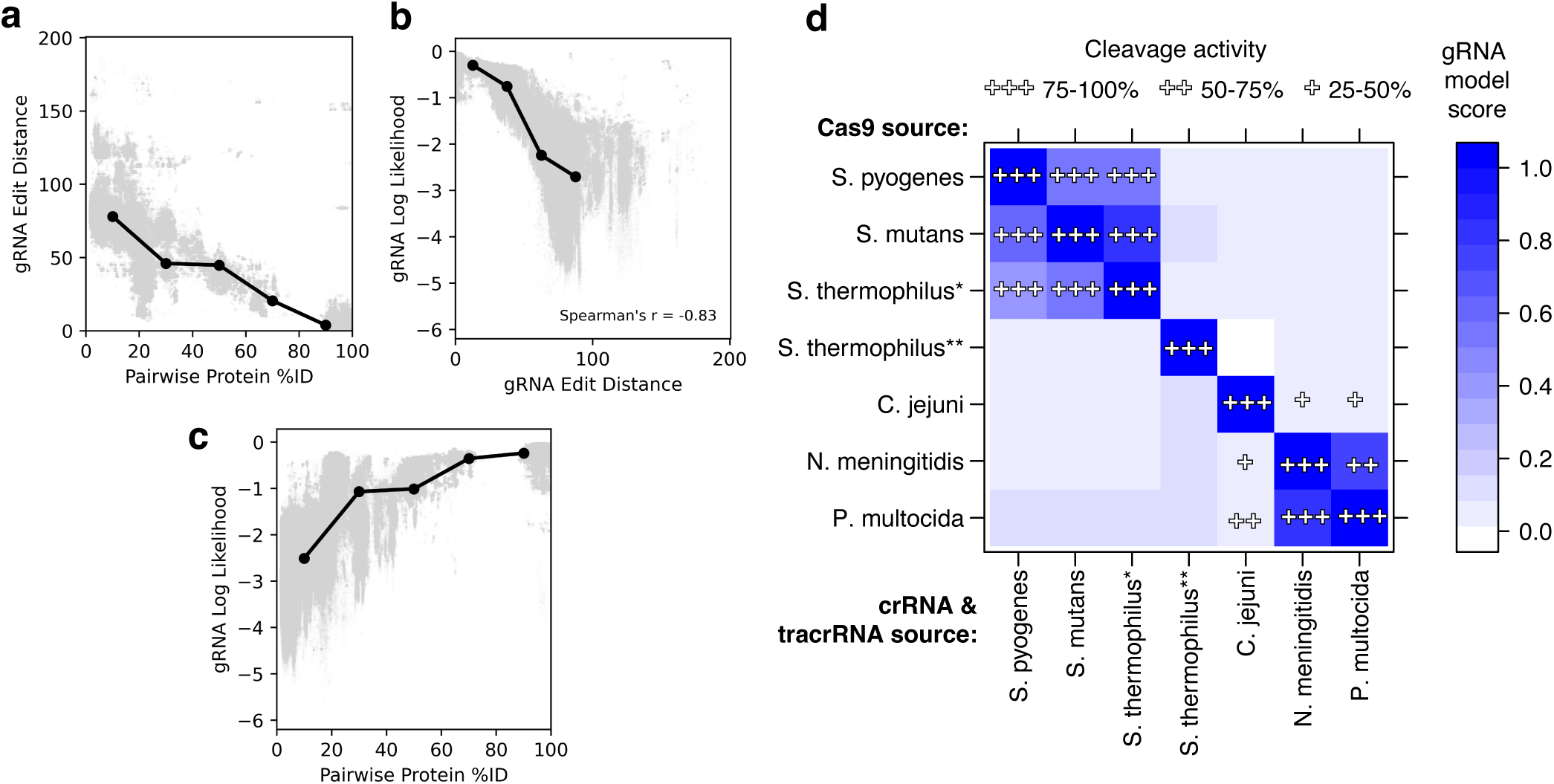
gRNA model predicts exchangeability of RNAs between orthologous Cas9s. a-c) crRNA:tracrRNA pairs were obtained for 1,591 distinct natural Cas9 sequences not used to train the gRNA model. The gRNA model was used to score native RNA:protein and non-native RNA:protein interactions. Additionally, pairwise identity was computed between RNA and protein sequences. a) RNA sequences diverge along with Cas9, but at a slower rate. b) RNA edit distance is correlated with the gRNA model score. c) gRNA model scores remain high for gRNAs exchanged between Cas9 proteins that display >70% identity. d) In 2013, Fonfara et al. tested the exchangeability of dual guide RNAs in vitro between eight diverse Type II CRISPR-cas systems (81). DNA cleavage rates from Fonfara et al. are displayed in the figure as: +++: 75-100%, ++: 50-75%, +: 25-50%. We applied the gRNA model to score each pair of Cas9:RNA sequence pairs. The gRNA model outputs a log likelihood for each protein:RNA pairing. The matrix shows a relative gRNA compatibility score, quantified as the softmax over the RNA log likelihoods for each protein. Note: Fonfara et al. also tested F. novicida Cas9 and RNA molecules; however, because the tracrRNA for this and related proteins were not found in our training set, we excluded F. novicida from this analysis.

**Fig. S8.**
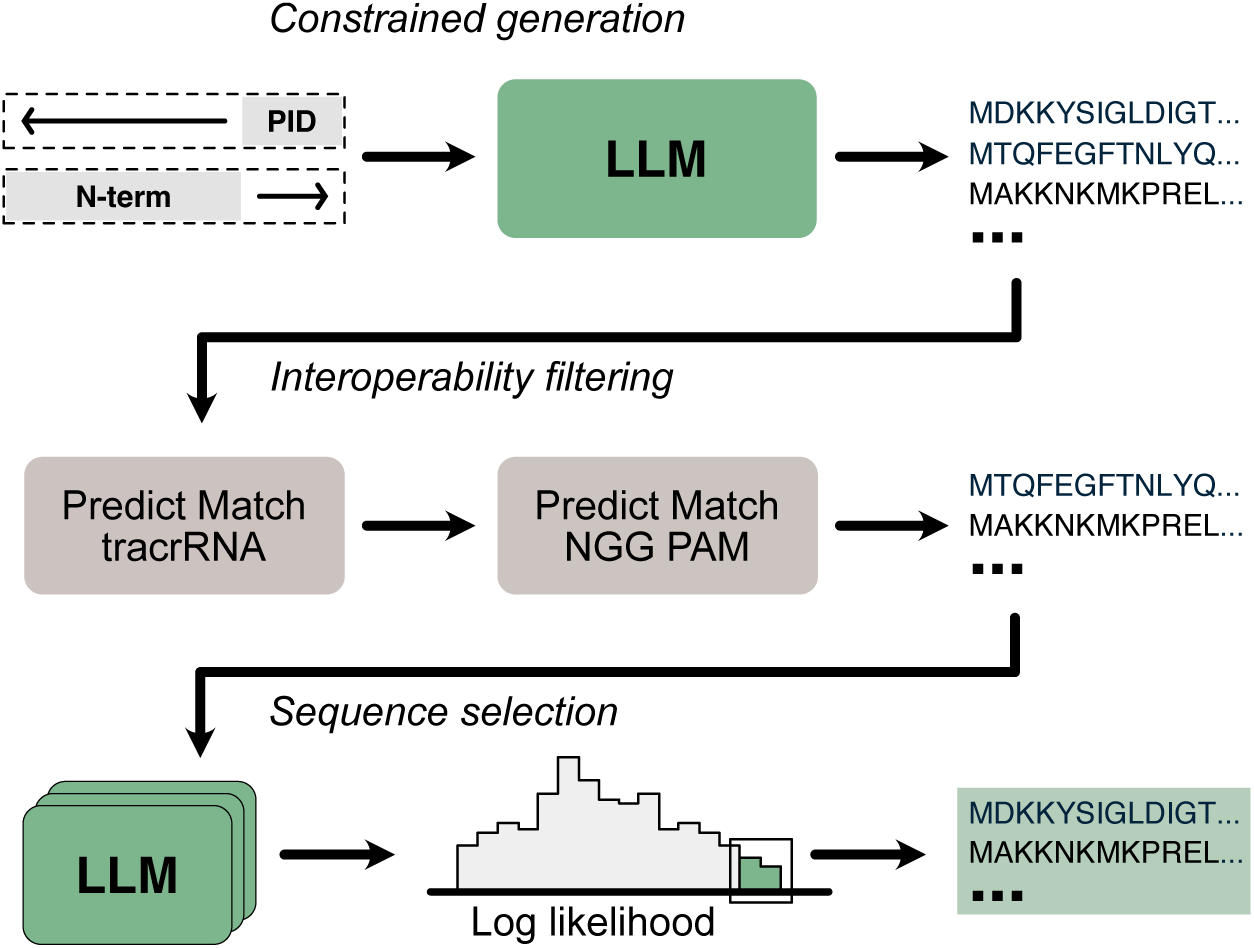
Constrained generation strategy for design of Cas9-like proteins. Sequences were generated conditionally based on the PID domain or N-terminal (non-PID) segments of SpCas9 (269 and 1,099 residues, respectively) using the Cas9 PLM. To select sequences for testing that would be compatible with SpCas9’s sgRNA and NGG PAM preference, we trained two sequence classifier models to use as interoperability filters. Sequences were ultimately selected for experimental characterization according to an ensemble of pre-trained and fine-tuned language model log likelihoods.

**Fig. S9.**
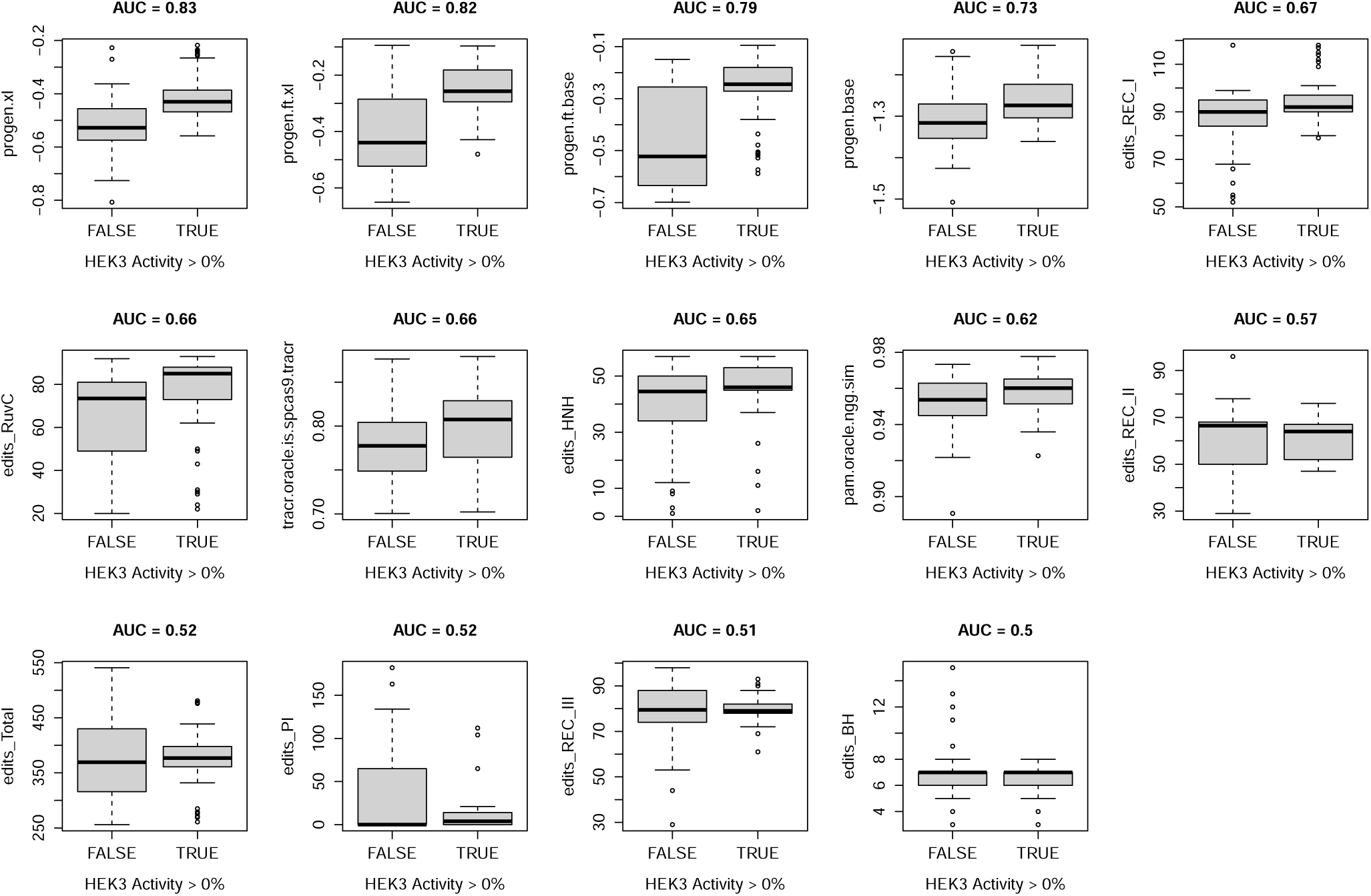
Language model scores predict enzyme activity. 209 Cas9-like proteins were screened for nuclease activity at the HEK3 target site. Boxplots indicate the distribution of various model scores and other statistics between active and inactive enzymes. Language model scores are indicated by y-axis labels: progen.xl, progen.ft.xl, progen.ft.base, and progen.base (key: xl=extra-large, ft=fine-tuned, base=base-model). Y-axis labels with prefix “edits_” indicate the Levenshtein distance to SpCas9 for a given domain. Y-axis labels “tracr.oracle.is.spcas9.tracer” and “pam.oracle.ngg.sim” indicate scores from models designed to determine whether a protein would be compatible with SpCas9’s tracrRNA or PAM.

**Fig. S10.**
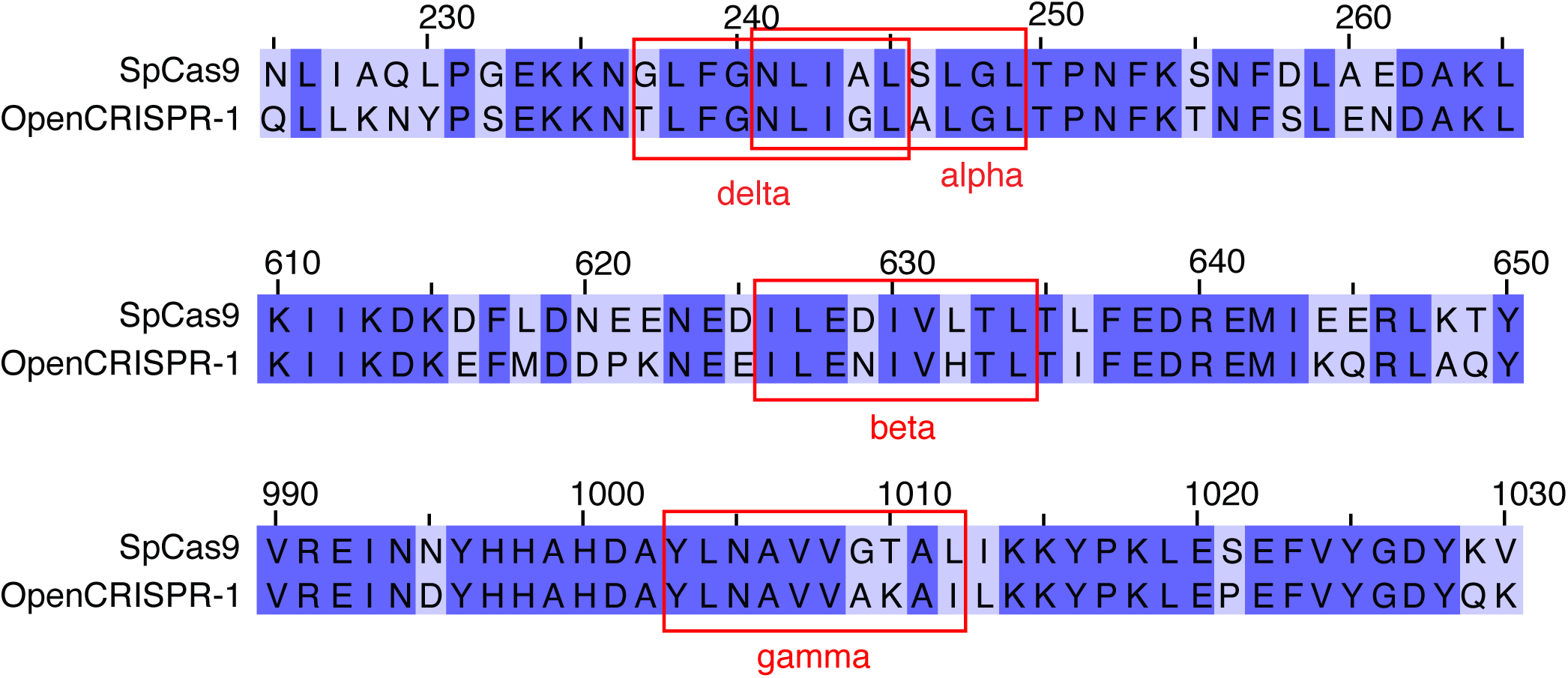
Comparison of SpCas9 immunogenic epitopes with OpenCRISPR-1. Ferdosi et al. discovered four SpCas9 epitopes that elicit pre-existing B cell and T cell immune responses in healthy donors (36). Alignments to SpCas9 and OpenCRISPR-1 revealed one or more mutations in each of these epitopes.

**Fig. S11.**
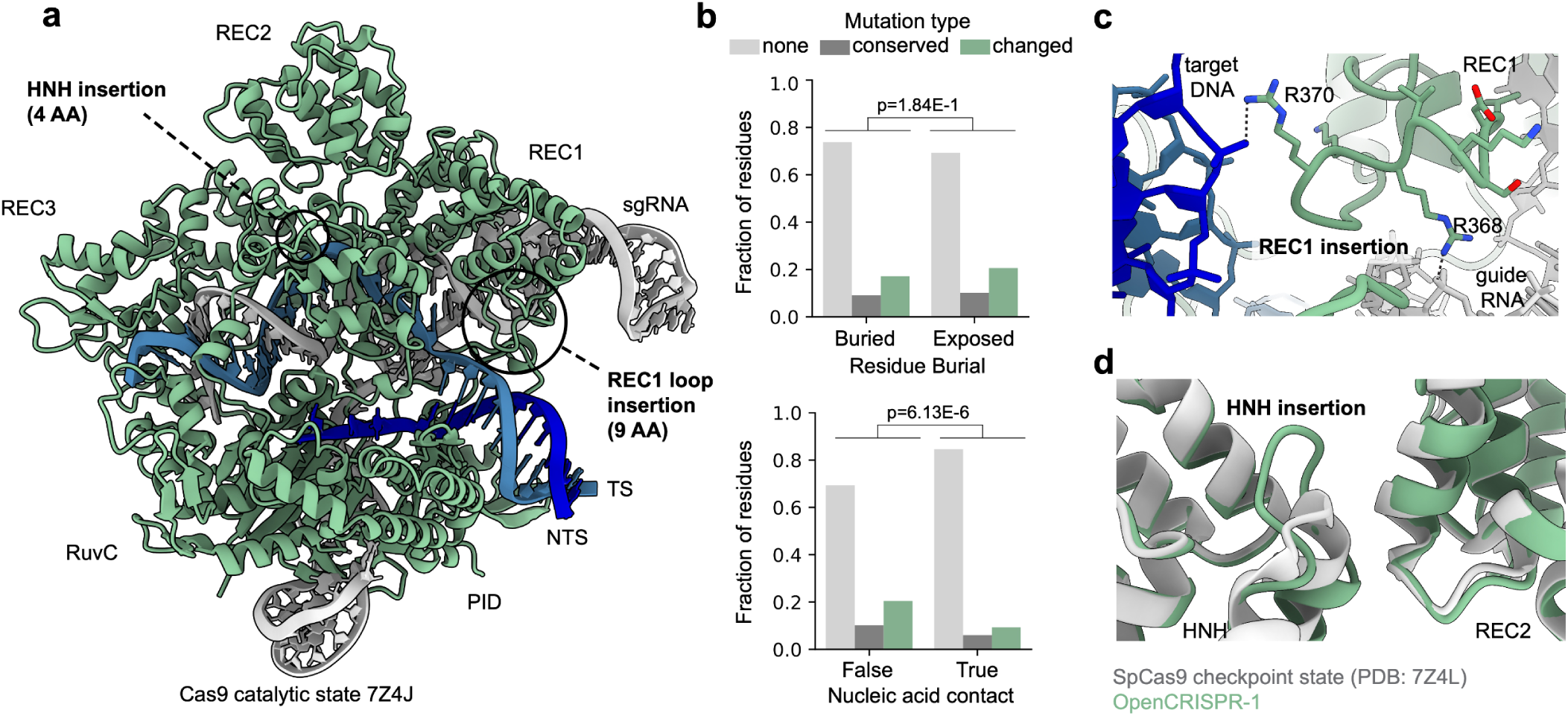
Structural analysis of OpenCRISPR-1. The structure of the OpenCRISPR-1 protein was predicted with AlphaFold2 (29) using multiple-sequence alignments from ColabFoldDB (79) and template structure of SpCas9’s catalytic state (PDB: 7Z4J). The predicted protein structure was aligned to complex with the sgRNA and DNA. a) Structural model of OpenCRISPR-1 effector complex in catalytic state. Insertions in the HNH and REC1 domains with potential functional implications are highlighted. b) Analysis of OpenCRISPR-1 mutational distribution relative to SpCas9 according to residue burial (top) and whether a residue is in contact (<4.0Å) with nucleic acids (bottom). Residue burial was not a significant determinant of mutational distribution, while nucleic-acid contacting residues were significantly depleted in mutations (chi-squared contingency test, p<0.05). c) Nine-residue positively charged insertion in the REC1 domain of OpenCRISPR-1, which introduces stabilizing interactions with the phosphate backbone of the guide RNA’s repeat:anti-repeat segment and the target DNA’s PAM-proximal region. d) Four-residue insertion in the HNH domain of OpenCRISPR-1 modeled in the checkpoint state (PDB: 7Z4L), which may serve to stabilize the cleavage checkpoint state.

**Fig. S12.**
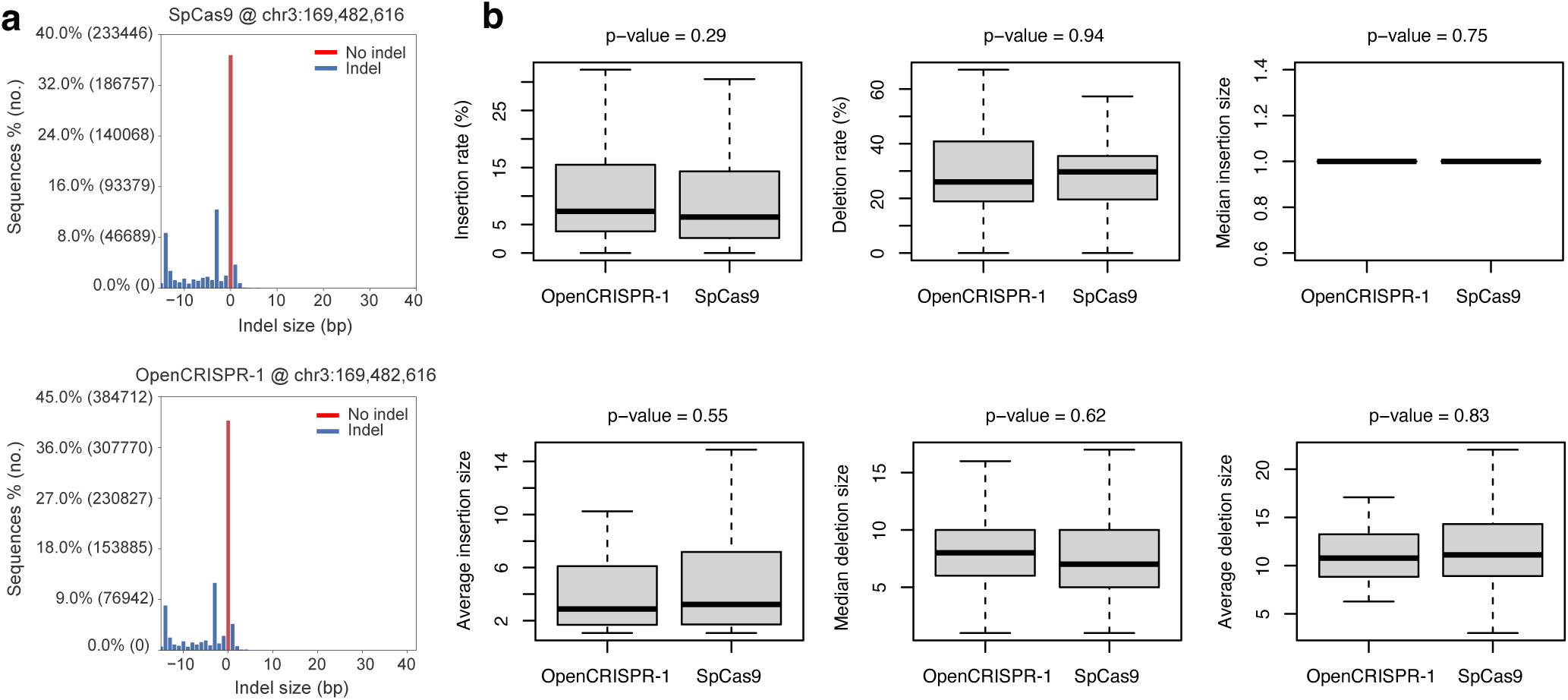
DNA repair outcomes for SpCas9 and OpenCRISPR-1. a) Comparison of the distribution of DNA repair outcomes between SpCas9 and OpenCRISPR-1 at one genomic site with CRISPResso2 (74). b) At each site, we summarized the distribution of insertions and deletions, and compared these statistics between proteins across 92 sites. Across the 92 sites, editing with SpCas9 and OpenCRISPR-1 result in similar DNA repair outcomes.

**Fig. S13.**
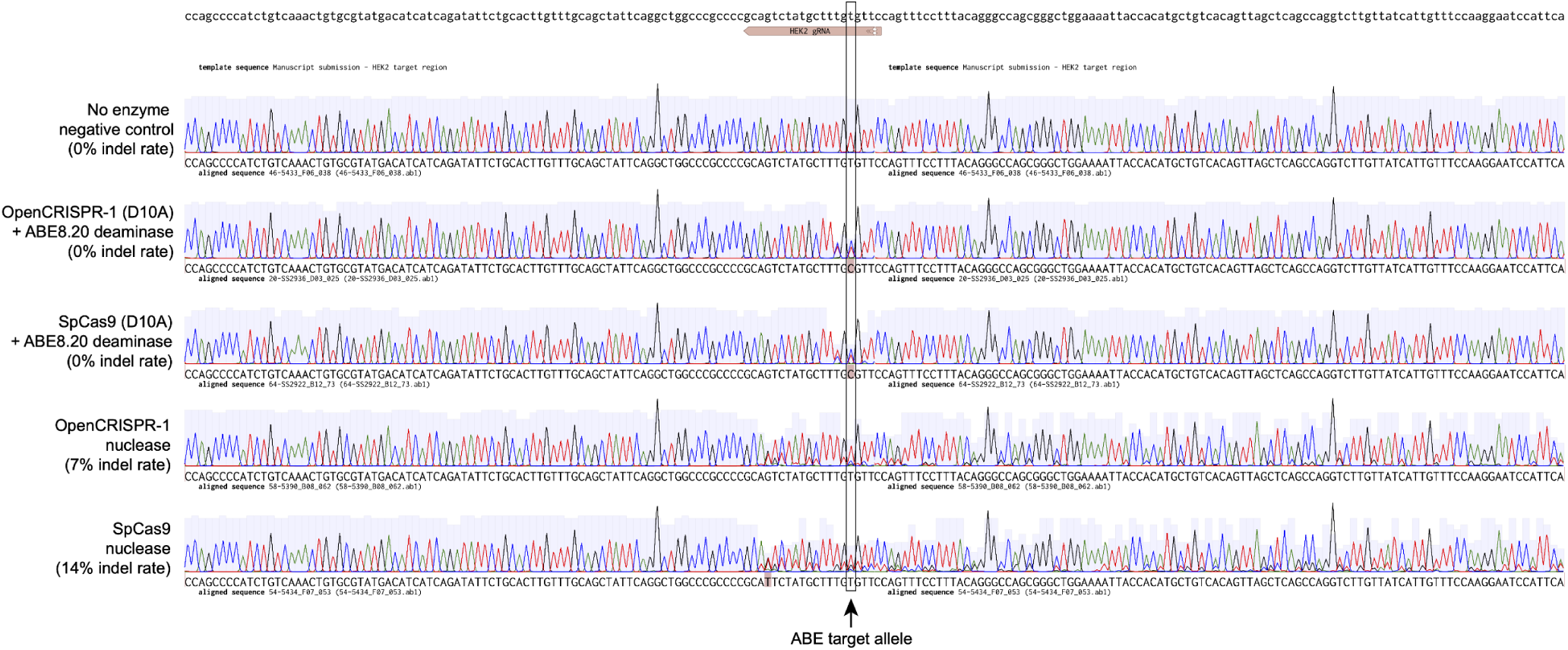
Converting OpenCRISPR-1 to a nickase via the D10A mutation. OpenCRISPR-1 was converted to a putative nickase via the D10A mutation. OpenCRISPR-1 and SpCas9 nuclease and nickases were used to target the HEK2 genomic site in HEK293T cells. Editing outcomes were assayed via Sanger sequencing. The plot above shows Sanger sequencing traces for a no enzyme negative control, OpenCRISPR-1-nickase ABE8.20 base editor, SpCas9-nickase ABE8.20 base editor, OpenCRISPR-1 nuclease, and SpCas9 nuclease. Indel rates were measured using the Inference of CRISPR Edits (ICE) program (31). As assayed via the ICE tool, the OpenCRISPR-1 nickase does not result in detectable double strand breaks.

**Fig. S14.**
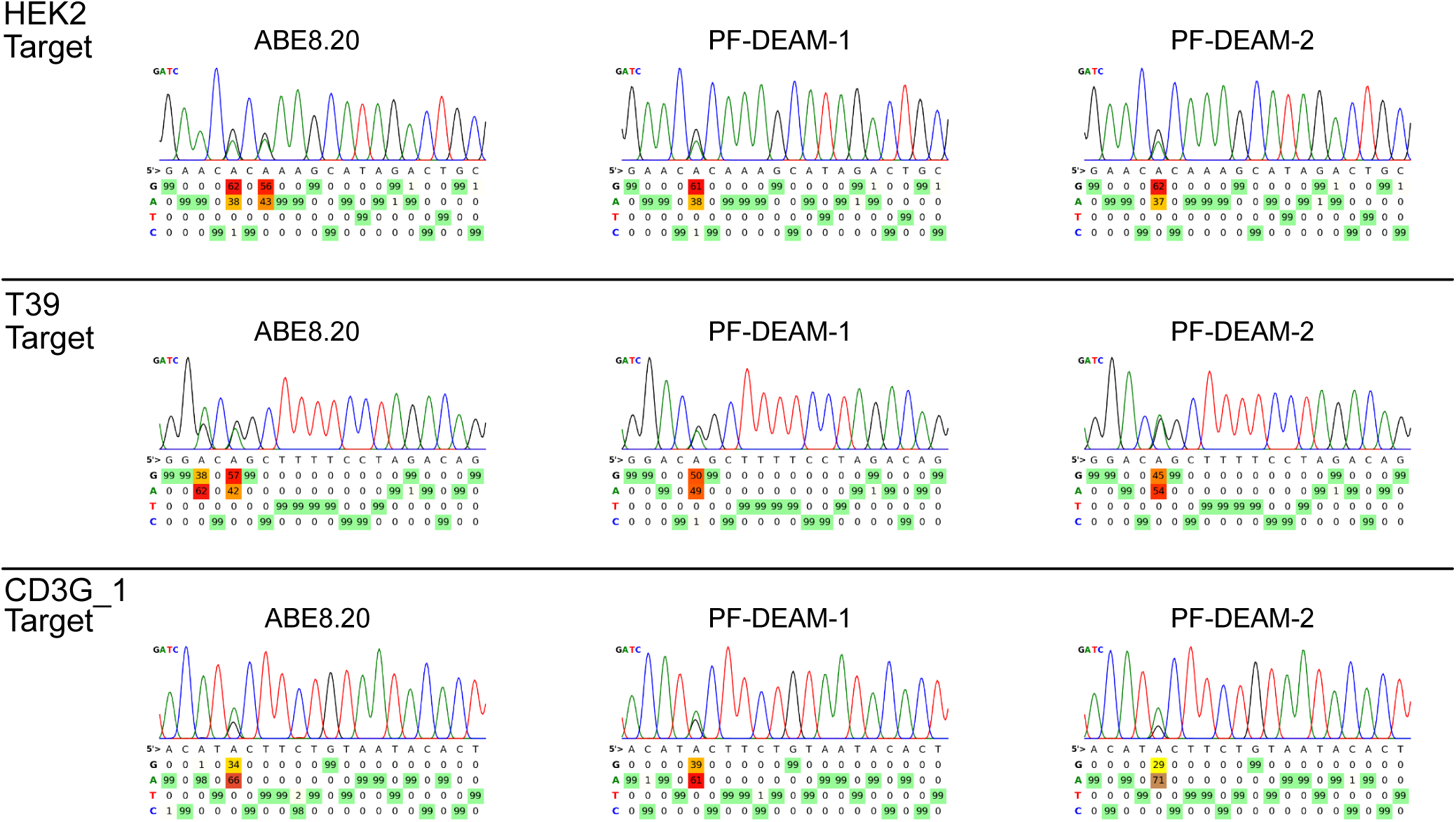
Base editing profiles of evolved and generated deaminases with OpenCRISPR-1. Deaminases were linked to an OpenCRISPR-1 nickase (D10A) using an N-terminal linker and tested in triplicate for base editing at three sites. All base editors displayed maximum A-to-G editing at position five, but constructs using generated deaminases produced significantly fewer (if any) bystander edits as compared to those using ABE8.20.

**Fig. S15.**
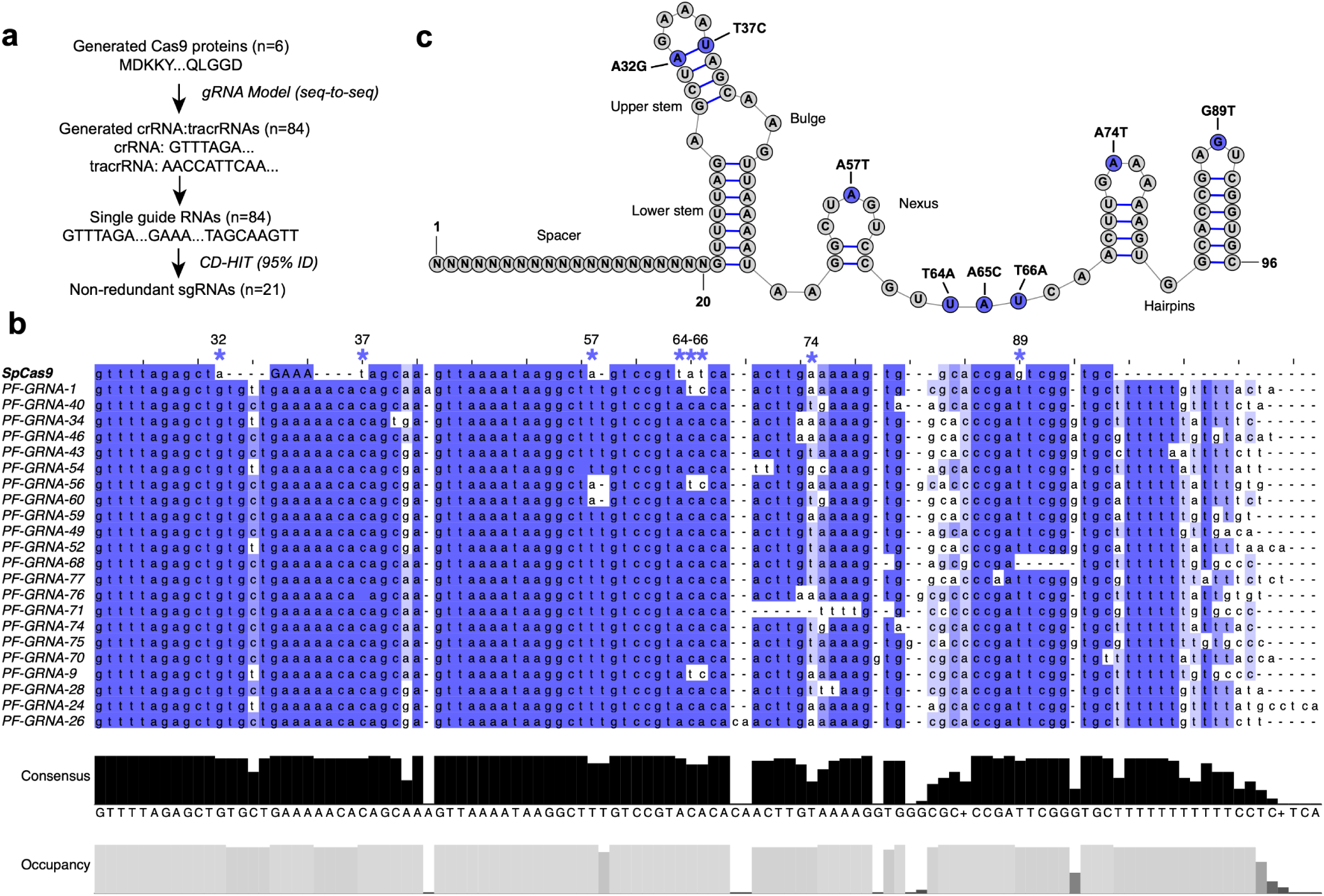
Comparison of generated and SpCas9 gRNA scaffolds. a) sgRNAs were generated for the most performant ML designed Cas9s. sgRNAs were de-replicated at 95% identity using cd-hit (64). b) Non-redundant sgRNAs were aligned using FAMSA (67) and visualized using Jalview (82). Blue stars indicate fixed, or nearly-fixed, differences between SpCas9 and generated sgRNA scaffolds. Terminator sequence for SpCas9 not shown. SpCas9 sgRNA is shown with a 12-nt stem loop, whereas generated sgRNAs were designed with a 16-nt stem loop. c) Predicted secondary structure of SpCas9s sgRNA with mutations overlaid.

**Fig. S16.**
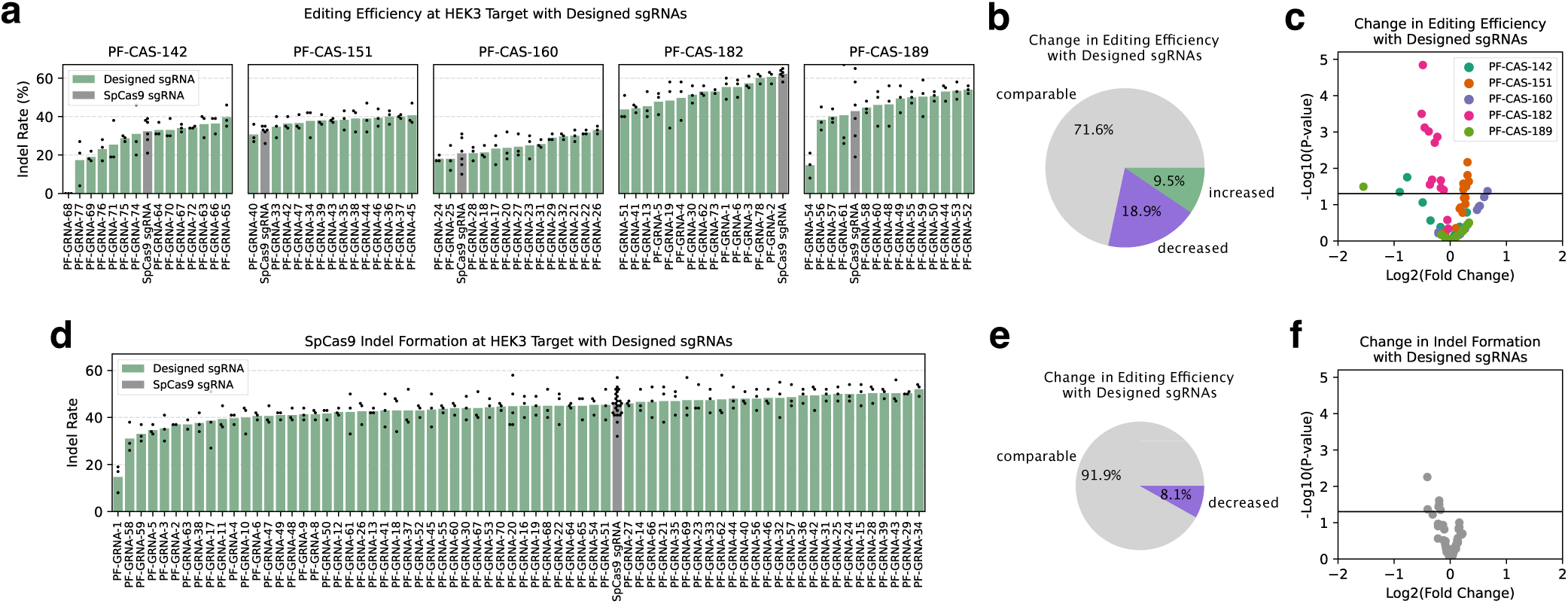
Editing efficiency with designed sgRNAs. Single-guide RNAs (sgRNAs) were designed for five generated nuclease proteins and tested for editing efficiency at the HEK3 target site with their cognate generated proteins and SpCas9. a) Editing efficiency with generated proteins and designed sgRNAs. b) Distribution of outcomes for designed sgRNAs paired with generated proteins. No change indicates insignificant change (P-value > 0.05) in editing efficiency with designed sgRNA for a given protein as compared to editing with SpCas9’s sgRNA. Improved and worsened correspond to significant increases and decreases in editing efficiency when a protein is paired with a designed sgRNA, respectively. c) Volcano plot showing significance of editing efficiency changes for each generated protein when paired with designed sgRNAs rather than SpCas9’s sgRNA. Horizontal line indicates a P-value of 0.05. d) Editing efficiency of SpCas9 with designed sgRNAs. e) Distribution of outcomes for designed sgRNAs paired with SpCas9. f) Volcano plot showing significance of editing efficiency changes SpCas9 when paired with designed sgRNAs rather than its naturally derived sgRNA.

